# Asynchronous mouse embryo polarization leads to heterogeneity in cell fate specification

**DOI:** 10.1101/2024.07.26.605266

**Authors:** Adiyant Lamba, Meng Zhu, Maciej Meglicki, Sylwia Czukiewska, Lakshmi Balasubramaniam, Ron Hadas, Nina Weishaupt, Ekta M. Patel, Yu Hua Kavanagh, Ran Wang, Naihe Jing, Magdalena Zernicka-Goetz

## Abstract

The first lineage allocation in mouse and human embryos separates the inner cell mass (ICM) from the outer trophectoderm (TE). This symmetry breaking event is executed through polarization of cells at the 8-cell stage and subsequent asymmetric divisions, generating polar (TE) and apolar (ICM) cells. Here, we show that embryo polarization is unexpectedly asynchronous. Cells polarizing at the early and late 8-cell stage have distinct molecular and morphological properties that direct their following lineage specification, with early polarizing cells being biased towards producing the TE lineage. More recent studies have also implicated heterogeneities between cells prior to the 8-cell stage in the first lineage allocation: cells exhibiting reduced methyltransferase CARM1 activity at the 4-cell stage are predisposed towards the TE fate. Here, we demonstrate that reduced CARM1 activity and upregulation of its substrate BAF155 promote early polarization and TE specification. These findings provide a link between asymmetries at the 4-cell stage and polarization at the 8-cell stage, mechanisms of the first lineage allocation that had been considered separate.

## Introduction

The mammalian embryo begins development as a totipotent zygote that can generate all embryonic and extra-embryonic cell lineages. Cells of the embryo progressively lose this totipotency and initiate the first differentiation event that separates the inner cell mass (ICM) lineage, which gives rise to the organism, from the extra-embryonic trophectoderm (TE) lineage that gives rise to the placenta.

This first lineage segregation of TE and ICM depends on polarization of the embryo at the late 8-cell stage (Johnson and Ziomek, 1980). During polarization, each blastomere gains a defined ‘apical domain’ through polarization of the actomyosin cytoskeleton to the cell-free contact area and then transport of PAR proteins to the cortex (Plusa et al., 2005; Vinot et al., 2005; Louvet et al., 1996; Zhu et al., 2017). After polarized blastomeres divide, either one or both daughter cells will retain this polarization status as the apical domain reassembles (Ziomek and Johnson 1982; Anani et al. 2014; Korotkevich et al. 2017; Lim et al. 2020). The polarized blastomeres will stay outside and form the TE, and inner blastomeres that do not have the apical domain will be specified as ICM. At the late 8-cell stage, after embryonic compaction, when cell-cell contacts increase and cells flatten against one another (Ducibella and Anderson, 1975; Zhu et al., 2017), all blastomeres are polarized.

Whether an 8-cell stage blastomere generates one inside and one outside daughter cell, or generates two outside daughter cells has been attributed to several factors, such as cell shape (Niwayama et al., 2019), nuclear positioning (Ajduk et al., 2014), transcription factor expression (Jedrusik et al., 2008) and spindle organization (Pomp et al., 2022). More recently, it has been shown that keratin filaments promote the inheritance of the apical domain as it reforms after the 8-16 cell stage division (Lim et al., 2020). Ultimately, cells with the apical domain will inactivate Hippo signalling, through sequestration of proteins such as LATS and AMOT, leading to YAP/TEAD4-mediated upregulation of CDX2 expression and specification as TE (Strumpf et al., 2005; Ralston and Rossant, 2008; Nishioka et al., 2008; Leung and Zernicka-Goetz, 2013). In contrast, apolar cells without the apical domain will have active Hippo signalling, lack nuclear YAP, and express pluripotency genes such as *Nanog* and *Oct4* to specify the ICM fate (Mitsui et al., 2003; Niwa et al., 2000)

The lineage segregation into ICM versus TE is also influenced by cell heterogeneities that exist before the 8-cell stage. Blastomeres of the 4-cell stage mouse embryo have been found to be unequal in their developmental fate and potential (Piotrowska-Nitsche et al., 2005; Bischoff et al., 2008; Tabansky et al., 2013). This developmental heterogeneity has been related to the heterogeneity in the activity of the arginine methyltransferase CARM1 (Torres-Padilla et al., 2007; Goolam et al., 2016; Parfitt and Zernicka-Goetz, 2010): blastomeres with the lowest CARM1 activity have the lowest methylation of CARM1 targets and contribute preferentially to TE, whereas cells with higher CARM1 becoming biased towards ICM (Torres-Padilla et al., 2007; Goolam et al., 2016; Parfitt and Zernicka-Goetz, 2010). CARM1 activity biases cells towards the ICM fate, at least in part because it promotes the expression of pluripotency genes via its methylation of H3R26 (Goolam et al., 2016; White et al., 2016; Plachta et al., 2011) and of BAF155, a subunit of the chromatin remodelling SWI/SNF complex (Schaniel et al., 2009; Lim et al., 2020; Panamarova et al., 2016). In agreement with this, it has been found that 4-cell blastomeres with lower CARM1 activity preferentially produce blastomeres with keratin filaments at the 8-cell stage that bias them to the TE fate (Lim et al., 2020). However, whether and how heterogeneities at the 4-cell stage relate to cell polarization at the 8-cell stage, has remained an open long-standing question.

Here, we show that the timing of blastomere polarization at the 8-cell stage is asynchronous, and that the heterogeneity between the blastomeres at the 4-cell stage influences cell fate specification through affecting the timing of blastomere polarization.

## Results

### Asynchrony in the timing of embryo polarization

To determine whether the timing of cell polarization affects subsequent embryo morphogenesis, we injected both blastomeres of 2-cell embryos with mRNA for *Ezrin-RFP* and performed time-lapse imaging to monitor formation of the apical domain in live embryos (Figure 1A). We used *Ezrin-RFP* as it labels the apical domain without disturbing development (Louvet et al., 1996; Zhu et al., 2017, 2020). We defined the dynamics of apical domain formation based on three criteria: length of the domain enriched for EZRIN-RFP; intensity of the EZRIN-RFP at the apical domain compared to the surrounding cell surface; and depth of the EZRIN-RFP enriched region (see *Materials and Methods*). We found that in most embryos, apical domains formed at the late 8-cell stage (Figure 1B) (Reeve and Ziomek, 1981; Zhu et al., 2017). However, unexpectedly, we noticed that some 8-cell stage blastomeres formed apical domains much earlier (Figure 1B). To explore this observation further, we analysed the number of blastomeres within the embryos in which we could detect early and late polarizing blastomeres. We were able to capture 19 out of 87 (21.8%) embryos that had blastomeres polarizing at the early rather than late 8-cell stage, within the first hour after its third cleavage division. When we plotted the fraction of blastomeres polarizing over time for 177 blastomeres, we observed a bimodal distribution with one peak at the early 8-cell stage (1 hour post 4-8 cell division) and a broader peak at later 8-cell stages (Figure 1C). Injection of the embryo with different concentration of *Ezrin-RFP* mRNA did not affect the frequency of blastomeres polarizing early (Figure 1-figure supplement 1A, B).

**Figure 1.**
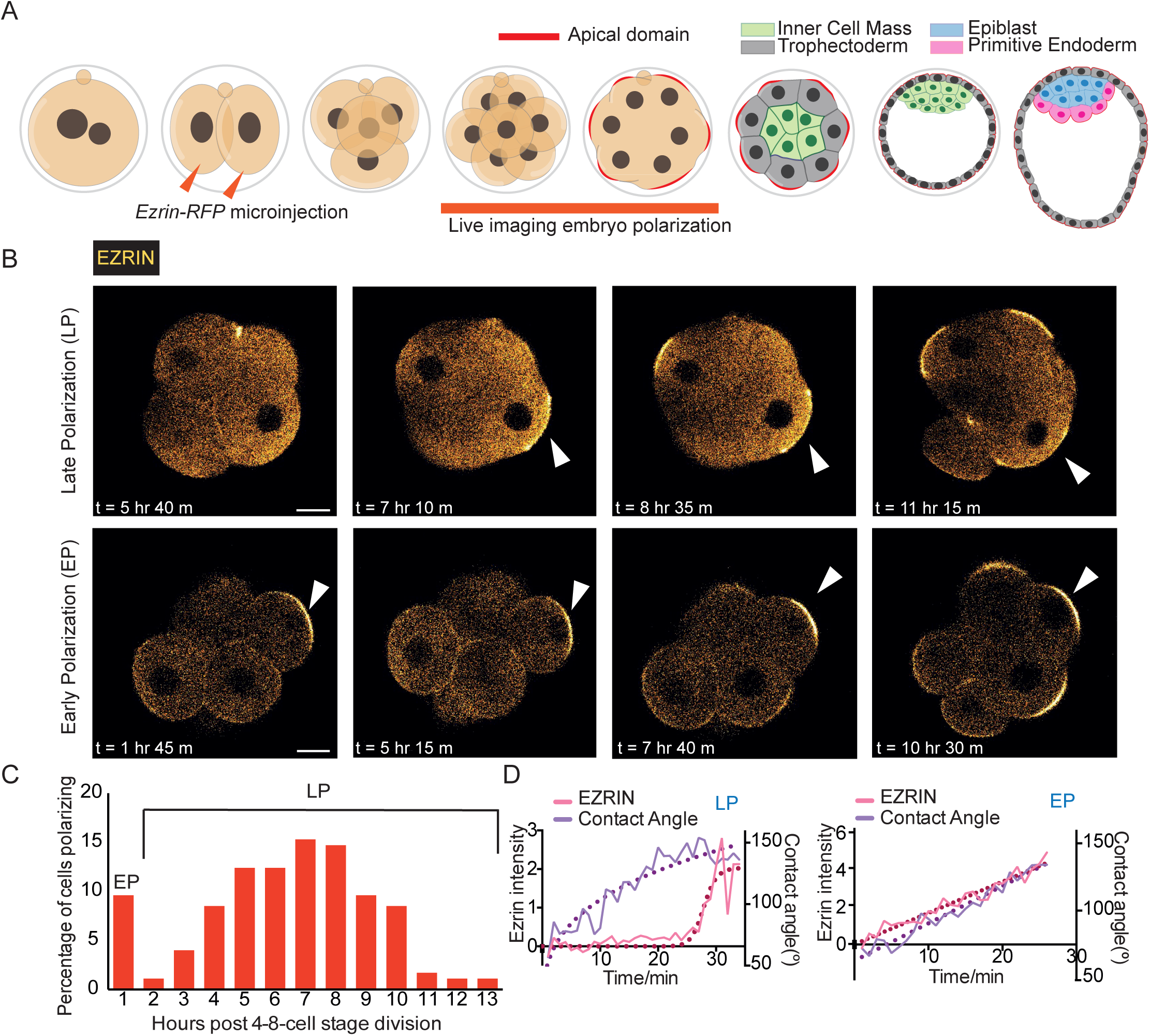
The timing of polarization at the 8-cell stage is asynchronous. A) Schematic representing mouse pre-implantation development and live imaging of apical-basal polarization through the 8-cell stage, with the polarization marker *Ezrin*-RFP microinjected at the 2-cell stage into both blastomeres. B) Time-lapse images of mouse embryos through the 8-cell stage showing EZRIN-RFP fluorescence, with apical domains indicated by the white arrowhead, and times post-8-cell stage division on each image. Apical domains (see *Materials and Methods* for definition) form at different times at the 8-cell stage. An example of a cell classed as ‘late polarization’ (LP) time in the top image and ‘early polarization’ (EP) time in the bottom image. C) Plot of polarization times throughout the mouse 8-cell stage, indicating two distinct peaks of polarization (‘early’ – EP, ‘late’ – LP), n = 177 blastomeres counted and polarization defined in *Methods*. D) EZRIN-RFP accumulation at the cortical surface and the inter-blastomere contact angle measured over time from the start of the changes in inter-blastomere angle associated with compaction, for representative LP and EP blastomeres – see *Materials and Methods* for details on calculation. Scale Bar = 15 µm.

To check whether this asynchronous polarization might be a result of live imaging of embryos, we also examined embryos that were fixed one hour post division to the 8-cell stage. We observed early polarizing blastomeres in 31.3% of fixed embryos, which is not significantly different than the frequency for live embryos (21.8%, Figure 1-figure supplement 1C). The proportion of embryos with early polarizing blastomeres was similar whether embryos were collected from super-ovulated mothers (21.8%) or from naturally ovulating mothers (25.0%) (Figure1-figure supplement 1D). Moreover, early polarizing blastomeres were observed at a similar frequency in different mouse strains (Figure 1-figure supplement 1E, F). Finally, the number of early polarizing blastomeres per embryo followed the theoretical number expected for an independent event, suggesting that polarization timing is a cell autonomous phenomenon (Figure 1-figure supplement 1G).

As the 8-cell stage progresses, the embryo compacts: the blastomeres pack tightly together, and the inter-blastomere contact angles increase from 60 to 180 degrees (Zhu et al., 2017). We therefore next compared the accumulation of apical EZRIN-RFP to the increase in the inter-blastomere contact angle (Zhu et al., 2017; Shen et al., 2022). For late polarizing blastomeres, the apical accumulation of EZRIN increased just before the plateau of the inter-blastomere contact angle, indicating that compaction begins before apical domain formation, as expected (Figure 1D). In contrast, in early polarizing blastomeres the increase in the inter-blastomere contact angle was concomitant with apical accumulation of EZRIN (Figure 1D). Thus, our data suggest that some blastomeres polarize before embryo compaction, while the majority polarize after the embryo compacts.

### Blastomeres polarizing early use the same mechanism as late polarizing blastomeres

As blastomeres polarize, F-ACTIN accumulates with EZRIN in the apical domain (Zhu et al., 2017), along with apical proteins such as PARD6 (Alarcon, 2010; Plusa et al., 2005; Vinot et al., 2005). We detected F-ACTIN and PARD6 in the apical domains of early polarizing blastomeres, similarly to late polarizing blastomeres (Figure 2A, B). The live imaging of embryos labelled with *Ezrin-RFP* allowed us to monitor polarization of blastomeres from their formation until the late 8-cell stage. Importantly, we noticed that early and late polarizing blastomeres had distinct features: the nucleus was closer to the apical domain and the apical domain was larger at the late 8-cell stage when they had polarized early (Figure 2C-E). These results suggest that the emergence of apical domain early correlates with its larger size and a closer proximity to the nucleus at the late 8-cell stage.

**Figure 2.**
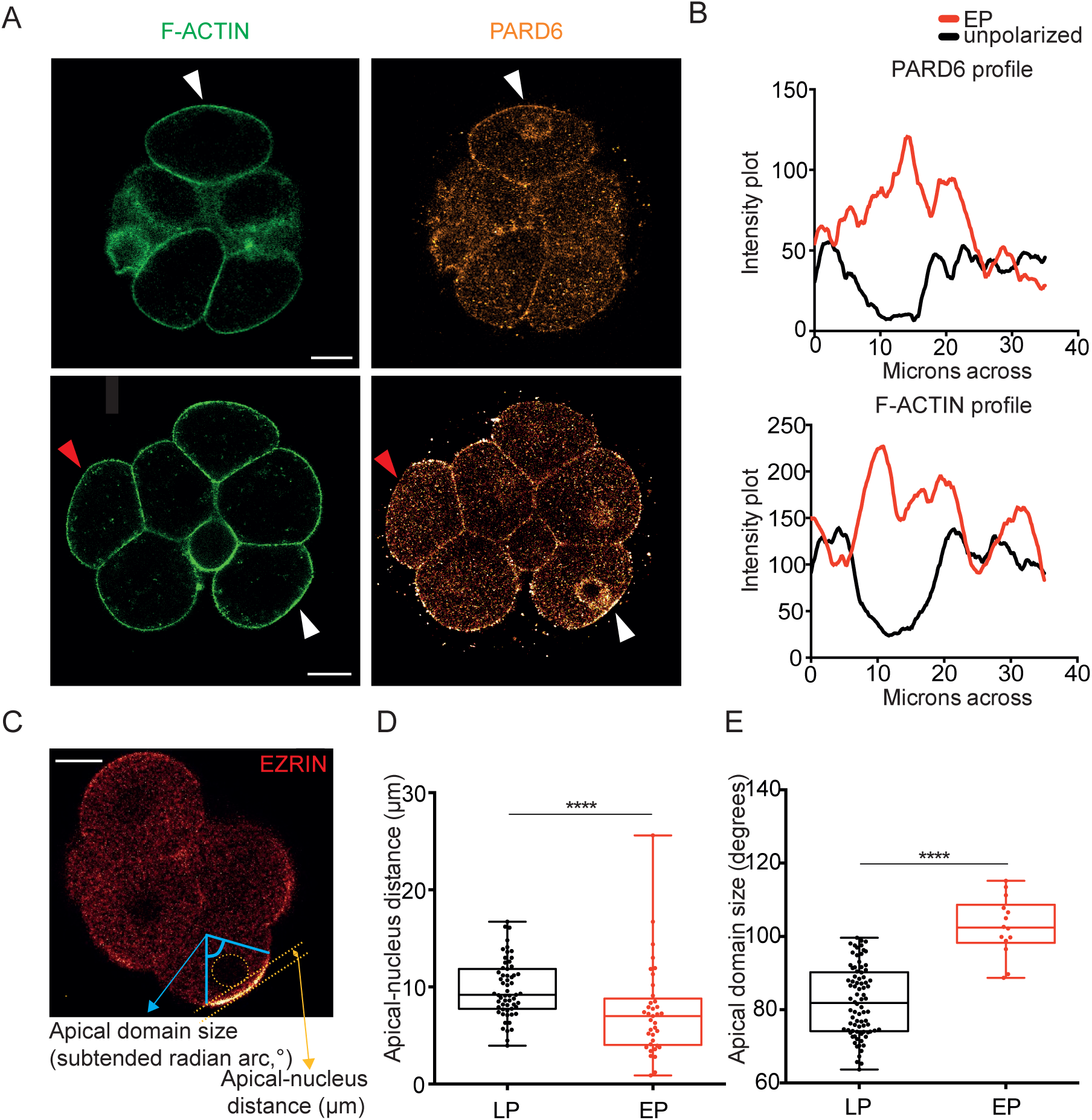
Early polarizing blastomeres accumulate PARD6 with F-ACTIN and have distinct morphological properties. A) Immunostaining of early 8-cell stage embryo, with white pointers indicting blastomeres containing both PARD6 and F-ACTIN accumulation at the outer cell cortex (EP) and red pointer indicating a blastomere with neither. B) Smoothened quantification of PARD6 and F-ACTIN across the outer membrane of an EP cell (white pointer, lower image) and an unpolarized cell (red pointer, lower image) at the early 8-cell stage. C) Method for calculating the apical-nucleus distance and apical domain size of blastomeres at the late 8-cell stage after assessment of polarity status, from live imaging using *Ezrin*-RFP, see *Materials and Methods* for further details. D) EP cells have a significantly smaller apical-nucleus distance than LP cells, Mann-Whitney test, ****p<0.0001, n = 22 EP cells, n = 59 LP cells analysed. E) EP cells have a significantly larger apical domain size than LP cells, two-tailed t test, ****p<0.0001, n = 14 EP cells, n = 82 LP cells. Scale Bar = 15 µm

Nuclear positioning was shown to be associated with microtubule organization (Laan et al. 2012a, 2012b; Dujardin and Vallee 2002). To establish whether microtubules are required for early blastomere polarization, we isolated single 4-cell stage blastomeres, as previously (Johnson and Ziomek, 1982.; Korotkevich et al., 2017), treated them with the microtubule depolymerizing drug colcemid (Siracusa et al., 1980) or vehicle (DMSO) and examined their division and the resulting “8-cell stage” doublets (Figure 2-figure supplement 1A). We found that the frequency of early polarization was unaltered by colcemid treatment, suggesting that microtubule polymerization is not required for early polarization (Figure 2-figure supplement 1B, C).

Polarization is known to require RHOA-mediated activation of the actomyosin network, and the transcription factors TFAP2C and TEAD4, which indirectly drive central clustering of apical proteins (Zhu et al., 2017, 2020). To test if these factors are involved also when blastomeres polarize early, we injected 2-cell stage blastomeres with dsRNAs targeting *Tead4* and *Tfap2c*, which abolishes their expression and polarization at the late 8-cell stage (Zhu et al., 2020), and used *Ezrin-RFP* to track polarization by live imaging (Figure 2-figure supplement 2A). We found that knockdown of *Tead4* and *Tfap2c* abolished polarization at the early 8-cell stage (Figure 2-figure supplement 2B). In addition, we treated 4-cell stage embryos with the RHOA inhibitor C3 Transferase, which abolishes polarization (Zhu et al. 2020), and found it also abolished polarization at the early 8-cell stage (Figure 2-figure supplement 2C-E). RHOA inhibition blocked cytokinesis but cells could still polarize (Zhu et al., 2020) (Figure 2-figure supplement 2D). These results indicate that early polarization, as late polarization, requires TFAP2C, TEAD4 and active RHOA.

### Blastomeres polarizing early upregulate expression of trophectoderm determinants

The apical domain inhibits Hippo signalling, driving nuclear translocation of YAP and increased expression of the TE lineage specifying transcription factor CDX2 (Nishioka et al., 2009, 2008). To assess these functions of the apical domain in early polarizing blastomeres, we first examined YAP localization in early 8-cell embryos. The levels of nuclear YAP were higher in the polarized blastomeres with an apical domain enriched for PARD6 than in unpolarized blastomeres lacking an apical domain (Figure 3-figure supplement 1A, B). In agreement with this, hybridization chain reaction (HCR) RNA fluorescence in situ hybridization of early 8-cell stage embryos revealed that the number of *Cdx2* mRNA puncta was higher in polarized blastomeres with a PARD6-positive apical domain than in unpolarized blastomeres, for 5 out of 6 embryos with EP cells (Figure 3A, B).

**Figure 3.**
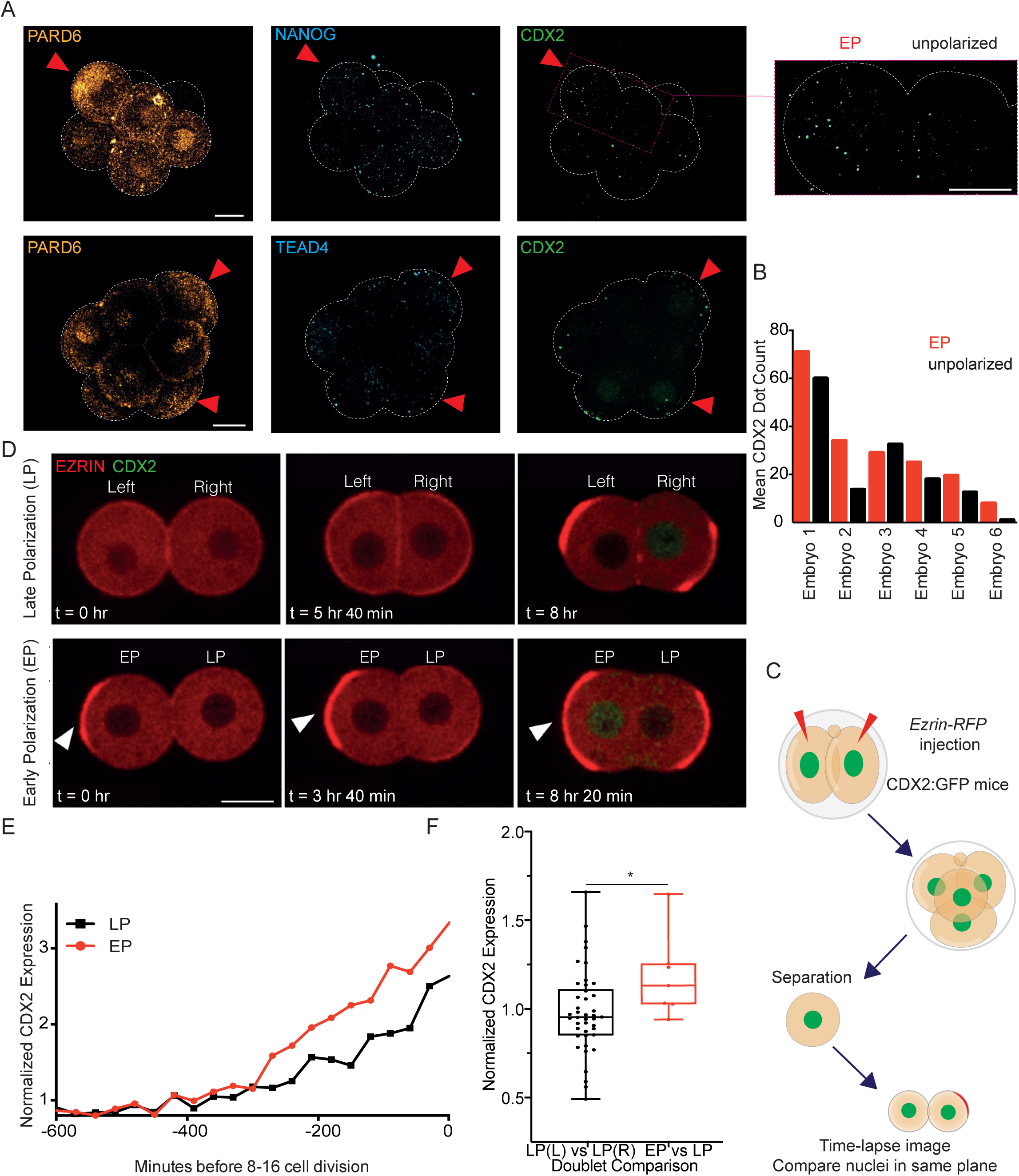
Early polarizing blastomeres show higher levels of *Cdx2* mRNA at the early 8-cell stage and CDX2 protein at the late 8-cell stage. A) Projection of representative Z slices in the uncompacted embryo. Immunostaining discerns EP (early polarizing) from unpolarized cells. Hybridization chain reaction for *Cdx2* and for *Nanog* or *Tead4* (positive controls, expression of these mRNAs is higher than *Cdx2* at the 8-cell stage in all cells (Deng et al., 2014). Red arrowhead indicates EP cells and the window shows an amplified display of *Cdx2* puncta. B) Mean number of *Cdx2* puncta in early polarizing (EP) and unpolarized cells. C) Schematic showing reduced system of 8-cell stage doublets. *Ezrin*-RFP mRNA is injected into both blastomeres of the CDX2:GFP 2-cell embryo; blastomeres are separated at the 4-cell stage and imaged until the late 8-cell stage. D) Representative images of doublets with nuclear CDX2-GFP and EZRIN-RFP. Top images show a doublet with two LP cells (‘late polarizing’), labelled as left and right, and the bottom shows a doublet with an EP and LP cell (‘early polarizing’). The white arrowhead indicated an EP cell. E) Normalized CDX2 intensity (to cytoplasmic background signal) over time in a representative doublet, ending at the first cell division that marks the final frame of the 8-cell stage. EP and LP cells are indicated. F) EP doublets have a significantly higher CDX2-GFP ratio when comparing the EP cell to the LP cell, than when comparing two LP cells in the LP doublets (Left to Right), Mann-Whitney test, n = 49 doublets total (7 EP), *p<0.05 Scale Bar = 15 µm.

Unlike *Cdx2* mRNA, CDX2 protein levels were not consistently elevated in early polarizing blastomeres versus the unpolarized blastomeres at the early 8-cell stage (Figure 3-figure supplement 1C, D). To determine whether this is due to a delay between the accumulation of *Cdx2* mRNA and CDX2 protein, we examined late 8-cell stage embryos. To this end, we injected *Ezrin*-RFP mRNA to the 2-cell stage transgenic embryos expressing a CDX2:GFP fusion protein from the *Cdx2* promoter (McDole and Zheng, 2012) and examined them at the late 8-cell stage. Since levels of CDX2-GFP between blastomeres were low, as an alternative method, we isolated blastomeres at the 4-cell stage and analyzed their development until they divide and form the “8-cell stage doublets”. This allowed us to compare nuclear CDX2-GFP intensity of all blastomeres in the same z-plane (Figure 3C-D). We compared levels of CDX2-GFP (normalized to the non-specific background detected in the cytoplasm) for each blastomere at the late 8-cell stage in seven doublets with an early polarizing cell and 42 doublets with both cells polarizing late (Figure 3E, F). We found that CDX2 ratios varied around a mean of 0.98 between cells that polarize late (when comparing the left cell to the right cell as a proxy for random arrangement), but around 1.18 between an early polarizing and late polarizing cell (Figure 3F), which is significantly higher. We infer that CDX2 levels vary between blastomeres, but that early polarizing cells have relatively high levels of CDX2 protein by the late 8-cell stage. These results suggest that early polarization leads to relatively higher levels of CDX2 at the last 8-cell stage.

### Blastomeres polarizing early are biased towards symmetric cell divisions and trophectoderm specification

Elevated levels of CDX2 have been shown to promote symmetric cell divisions at the 8-16 and 16-32 cell stages (Jedrusik et al., 2008). In addition, a more apically localized nucleus has been implicated in a bias towards symmetric cell divisions (Ajduk et al., 2014). We therefore considered that blastomeres polarizing early might show an increased frequency of symmetric divisions compared to late polarizing blastomeres as they show both elevated CDX2 expression and a more apically localized nucleus. To test this, we performed live imaging of 2-cell embryos injected with *Ezrin-RFP* mRNA throughout the 8-cell stage and up until the blastocyst stage (E3.5) (Figure 4A-B). This approach allowed us to identify early polarizing blastomeres, that form before compaction, and late polarizing blastomeres that form after compaction, and follow them until their lineage specification.

**Figure 4.**
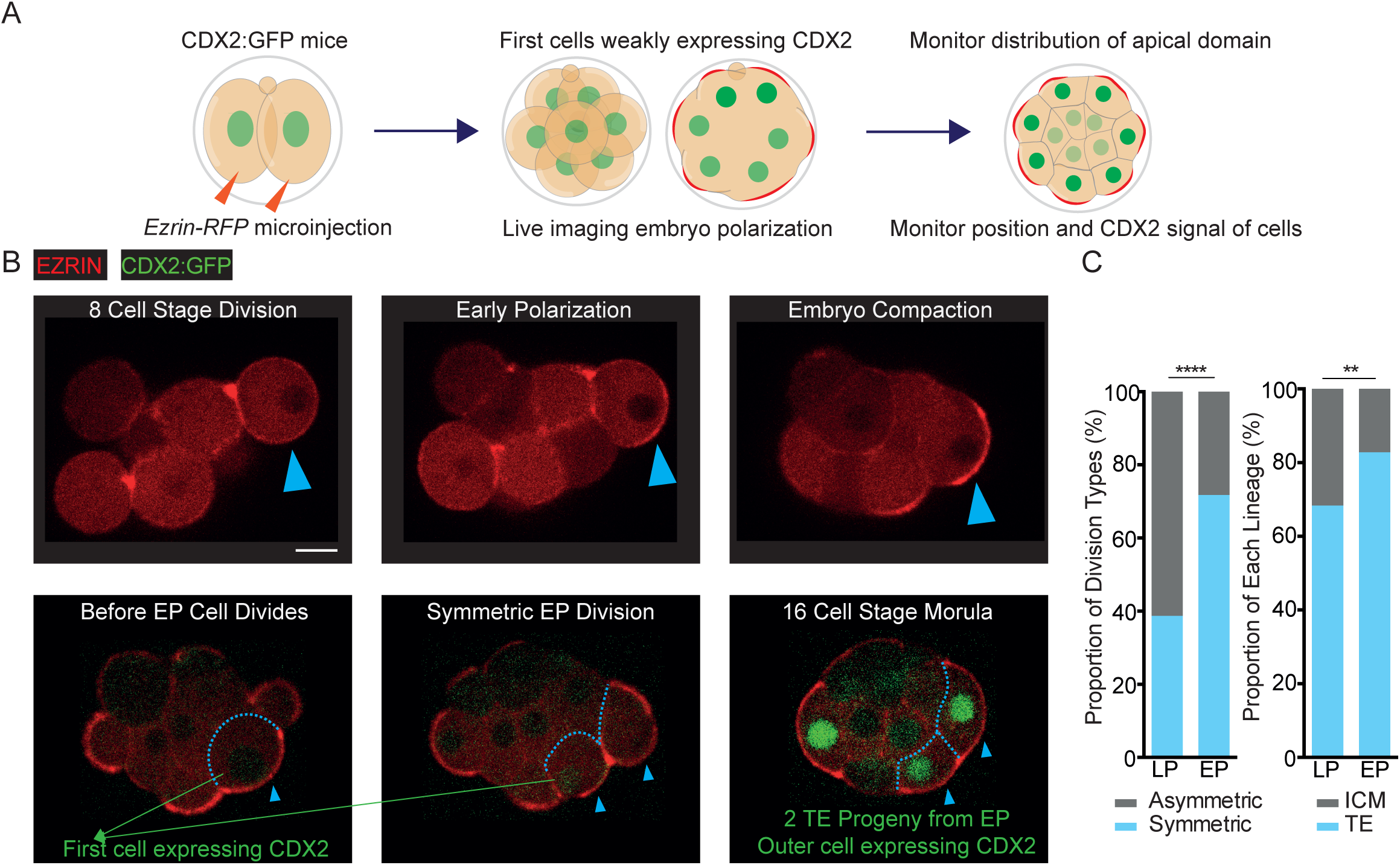
Early polarizing blastomeres are biased towards symmetric cell divisions and the trophectoderm lineage. A) Schematic indicating injection and time-lapse imaging to assess apical domain formation via EZRIN-RFP, and nuclear CDX2-GFP accumulation. B) Representative images showing time-lapse recordings of embryos over pre-implantation development during and after the 8-cell stage. Apical domain formation throughout the 8-cell stage is monitored by following EZRIN-RFP. Symmetric divisions are those in which both daughter cells inherit the EZRIN-RFP-positive apical domain after cell division, whereas in asymmetric divisions only one cell inherits the apical domain. Trophectoderm contribution can be assessed using the CDX2:GFP signal – outer, CDX2-GFP positive cells are counted as trophectoderm. C) EP cells have significantly greater frequency of symmetric divisions (left) and trophectoderm progeny (right). Two-tailed Fischer’s exact tests, ****p < 0.0001 and **p < 0.01 respectively; n = 199 LP cells and 46 EP cells were counted from, with 77 symmetric LP divisions (38.7%) and 33 EP symmetric divisions (71.7%); n = 275 LP cells and n = 87 EP cell progeny analyzed for lineage contribution at the 16-cell stage, with 188 LP trophectoderm cells (68.3%) and n = 72 EP trophectoderm cells (82.8%). Scale Bar = 15 μm.

To decipher asymmetric versus symmetric divisions, we followed the distribution of the EZRIN-RFP signal (Figure 4B). We found that 33 out of 46 early polarizing blastomeres (71.7%) divided symmetrically to generate 66 polarized blastomeres with an EZRIN-RFP-positive apical domain at the 16-cell stage (Figure 4C). The remaining 13 early polarizing blastomeres divided asymmetrically, giving rise to 13 polarized cells with an EZRIN-RFP-positive apical domain (79 polarized cells total, 85.9%) and 13 non-polar cells lacking an apical domain at the 16-cell stage (Figure 4C). In drastic contrast, only 77 out of 199 late polarizing blastomeres (38.5%) divided symmetrically and the remaining 122 late polarizing blastomeres divided asymmetrically, giving rise to 276 total polarized cells (69.3%) and 122 non-polar cells (Figure 4C).

We found that 72 out of 87 16-cell stage cells arising from early polarizing blastomeres contributed to the TE lineage (82.8%), whereas only 188 of 275 16-cell stage cells arising from late polarizing blastomeres gave rise to TE (68.3%; Figure 4C). These results indicate that the increased symmetric divisions of early polarizing blastomeres leads to their fate bias towards the TE lineage.

### Early polarization is associated with a change in cell geometry

The geometry of a polarized cell was shown to influence its decision to divide symmetrically or asymmetrically at the 8-16 cell stage (Niwayama et al., 2019; Korotkevich et al., 2017). Specifically, a higher width-to-height ratio (wider cells) can increase the proportion of symmetric cell divisions by orienting the spindle parallel to the axis of the apical domain, instead of its natural orientation towards the apical domain. To investigate whether the higher proportion of symmetric cell divisions in earlier versus later polarizing blastomeres reflects differences in shape, we carried out a 2-dimensional analysis of cell shape at the 8-16 cell division in the mid-plane of the embryo in time-lapse movies. *Ezrin*-RFP and *Life Act*-GFP mRNAs were co-injected at the 2-cell stage to follow the formation and presence of the apical domain and the cell boundaries, respectively (Figure 5A). Consistent with previous work, wider cells with an overall higher side: full length ratio and larger apex angle gave rise to more symmetric cell divisions, on average (Niwayama et al., 2019) (Figure 5B). Moreover, early polarizing blastomeres had wider morphology than late polarizing blastomeres (Figure 5C).

**Figure 5.**
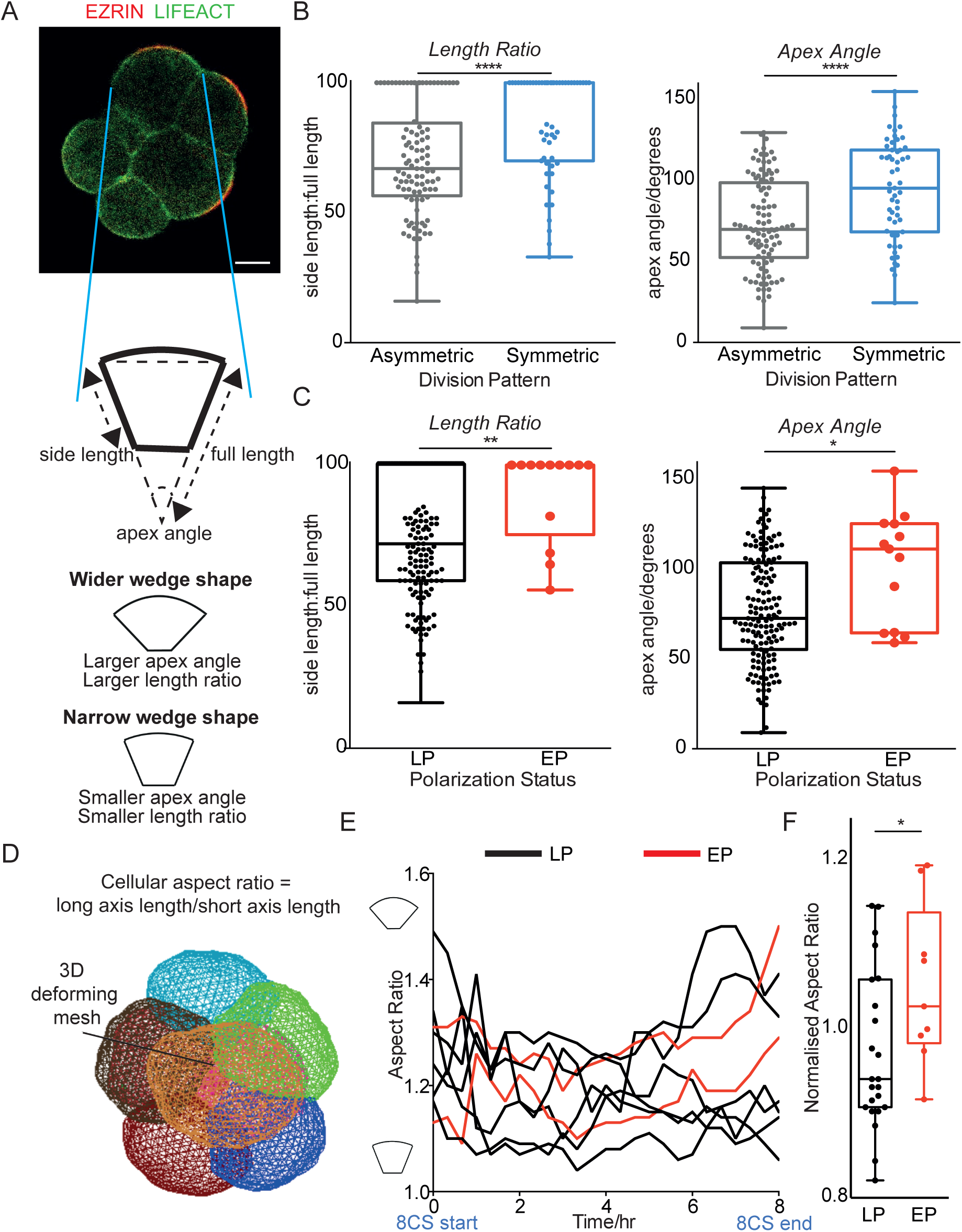
Early polarizing blastomeres have increased blastomere width. A) Indication of method for measuring the apex angle (degrees), side length and full length (*μ*m) in the embryo mid-plane before cell division, using time-lapse images with EZRIN-RFP and LIFEACT-GFP markers. B) Blastomeres dividing symmetrically have a higher length ratio and larger apex angle than asymmetrically dividing blastomeres. n = 45 embryos with n = 100 asymmetrically dividing and n = 55 symmetrically dividing cells analyzed. Any embryos with ambiguous division pattern were excluded, two-tailed student’s t-test, ****p < 0.0001 in both tests. C) Blastomeres from B re-classified according to their polarity status, with 13 EP blastomeres and 155 LP blastomeres identified. Early polarizing blastomeres have a higher length ratio and larger apex angle than LP blastomeres, two-tailed t-test, **p < 0.01 and *p < 0.05 respectively. Any embryos with ambiguous polarity status were excluded. D) Snapshot of 3D analysis of 8-cell embryos using 3D Mesh Deformation Plugin on Fiji – see *Method* for further details. E) Example embryo with EP and LP cells indicated, with their aspect ratio over time. A higher aspect ratio indicates a higher ratio of long axis: short axis length. F) 32 cells from 4 embryos with at least 2 EP cells each were analyzed in 3D at the end of the 8-cell stage, and aspect ratio measured and normalized to the embryo average. EP cells have a significantly larger normalized aspect ratio than LP cells, two-tailed t-test, *p < 0.05. Scale Bar = 15 µm.

To better visualize cell shape, we performed a 3D analysis on embryos with two or more early polarizing cells (Figure 5D, see *Materials and Methods)* to determine the ratio of the long axis to the short axis, which is larger in wider cells (Figure 5D). Plotting this cellular aspect ratio over time showed that early polarizing cells acquired a larger aspect ratio than late polarizing cells as the 8-cell stage progressed, within the same embryo (Figure 5E). Early polarizing blastomeres had a significantly wider aspect ratio before division than late polarizing cells, which is consistent with our 2D findings (Figure 5F). Thus, we infer that early polarizing cells have a wider shape than later polarizing cells at the 8-16 cell stage division, which is consistent with a bias towards symmetric division and the TE lineage.

### Asymmetries at the 4-cell stage influence the timing of cell polarization

We next wished to determine what causes some blastomeres to polarize earlier than others at the 8-cell stage. Initially, we hypothesised that differences in the timing of cell division at the 4-8 cell stage are responsible for differential polarization timing. Using cell tracking with *Ezrin-RFP*, we found that neither the time of the third cleavage division (4-to-8 cell stage), nor the division order at the 4-to-8 cell division, significantly correlated with early polarization (Figure 6-figure supplement 1).

We therefore wondered whether the asymmetries in the timing of cell polarization might relate to earlier asymmetries reported at the 4-cell stage (Piotrowska-Nitsche et al., 2005; Torres-Padilla et al., 2007; White et al., 2016). We previously found that blastomeres with low CARM1 activity in the 4-cell embryo are biased towards higher levels of CDX2 expression, symmetric cell divisions and the TE lineage (Torres-Padilla et al., 2007; Parfitt and Zernicka-Goetz, 2010). Given that early polarizing cells are also biased towards expressing higher levels of CDX2, symmetric cell divisions and the TE lineage, we hypothesized that lower levels of CARM1 would promote early polarization.

To test this possibility, we inhibited CARM1 activity in individual blastomeres, as previously (Panamarova et al., 2016; Parfitt and Zernicka-Goetz, 2010; Torres-Padilla et al., 2007), and determined if this manipulation can promote early polarization. We injected one blastomere at the 2-cell stage with mRNAs encoding *Ezrin-RFP* and catalytically inactive *HA-tagged CARM1(E267Q)*, or with *Ezrin-RFP* mRNA only (Figure 6A). CARM1 di-methylates H3R26 and we found that blastomeres with CARM1(E267Q) had a significant decrease in nuclear H3R26me2 intensity, as expected (Figure 6-figure supplement 2A, B). Strikingly, we observed a higher proportion of early polarizing blastomeres in the early 8-cell stage embryos co-expressing CARM1(E267Q) and EZRIN-RFP compared to those expressing only EZRIN-RFP (Figure 6B, C). As an orthogonal approach, we treated 2-cell stage embryos with a CARM1 inhibitor, *bis*-benzylidene piperidinone (Cheng et al., 2011), until the early 8-cell stage. We found that this treatment also caused reduced levels of H3R26me2 compared to control blastomeres (Figure 6-figure supplement 2C, D). Importantly embryos treated with the CARM1 inhibitor also showed a greater proportion of early polarizing cells than control-treated embryos (Figure 6C). Thus, decreased CARM1 activity increases the frequency of early polarizing cells.

**Figure 6.**
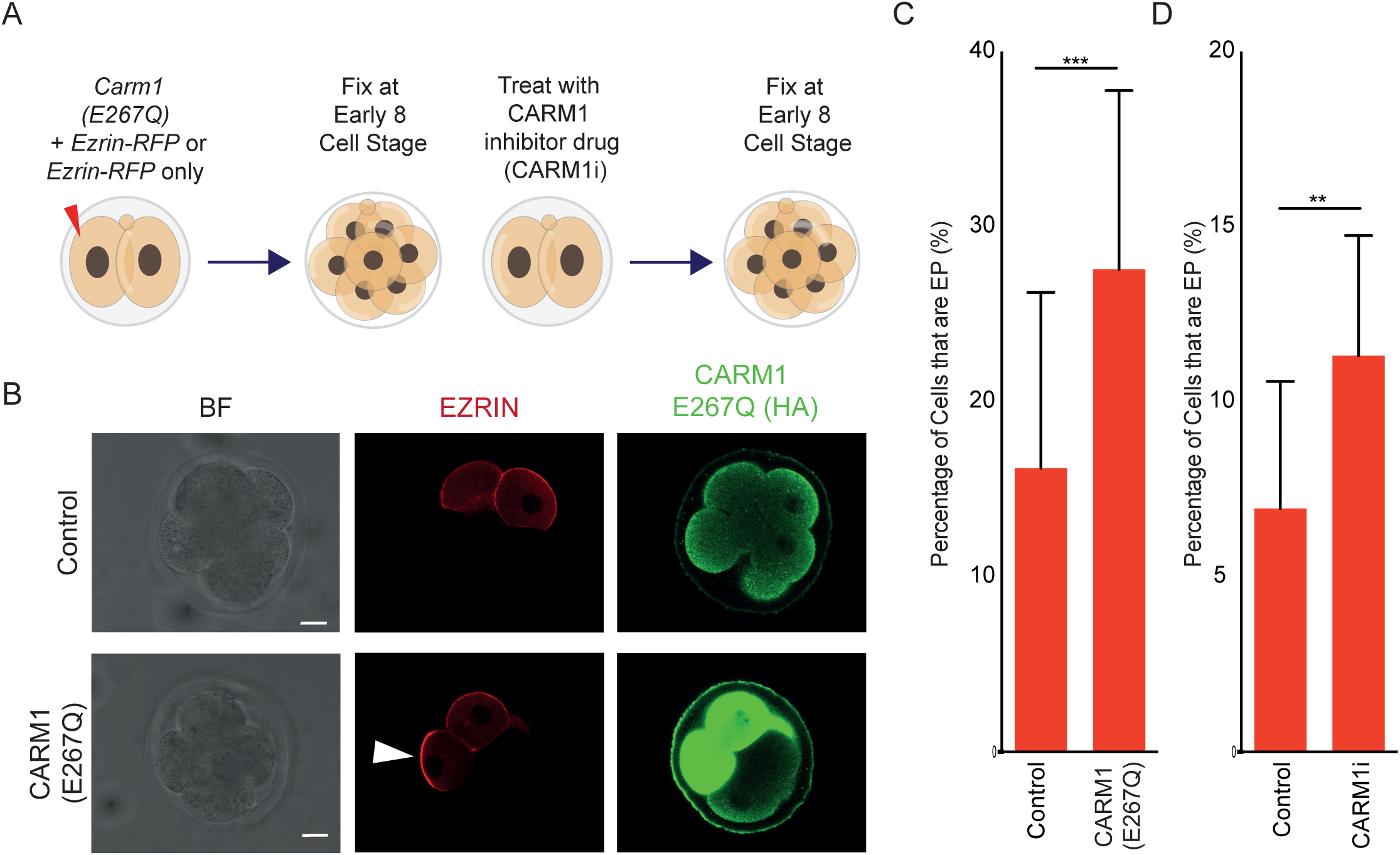
Frequency of early polarization is increased after inhibition of CARM1 activity. A) Schematic indicating experimental design for CARM1 inhibition using CARM1(E267Q) or CARM1 inhibitor. B) Immunostaining for DAPI, HA-tag (tagged to the CARM1(E267Q) construct), and injected *Ezrin*-RFP (at the early 8-cell stage). EZRIN-RFP present in injected blastomeres and an EP blastomere is indicated via an arrow. C) CARM1(E267Q) blastomeres have a significantly higher frequency of being early polarized than Control (*Ezrin*-RFP only) injected blastomeres at the early 8-cell stage, N = 4 independent experiments, two-tailed z-test, **p < 0.01; overall n = 97 CARM1 (E267Q) embryos with n = 385 cells (73 EP, 19.0%), n = 100 Control embryos with n = 377 cells (37 EP, 9.8%). Mean of individual experiments with S.E.M given on the graph. D) CARM1i blastomeres have a significantly higher frequency of being early polarized than Control (*Ezrin*-RFP only) injected blastomeres at the early 8-cell stage, two-tailed z-test, ***p < 0.001; N = 2 independent experiments, overall n = 32 CARM1i embryos with n = 372 cells (40 EP, 11%), n = 47 Control embryos with n = 376 cells (20 EP, 5 %). Mean of individual experiments with S.E.M given on the graph. Scale Bar = 15 µm.

In addition to H3, the SWI/SNF component BAF155 is also a substrate of CARM1. A decreased level of CARM1-mediated BAF155 methylation results in BAF complex stability, and subsequently its suppression of pluripotency genes (Panamarova et al., 2016). We and others have shown that BAF155 overexpression in zygotes leads to an increase in the number of CDX2-expressing (TE) cells at the blastocyst (Panamarova et al., 2016; Lim et al., 2020). To investigate if BAF155 overexpression, can change the frequency of early polarization, we co-injected one blastomere at the 2-cell stage with mRNAs encoding *Ezrin -RFP* and an *HA-tagged BAF155* construct, as previously (Panamarova et al., 2016) (Figure 7A). We observed an increased frequency of early polarizing cells in BAF155-overexpressing embryos compared with those expressing EZRIN-RFP only (Figure 7B, C), indicating that elevated levels of BAF155 can also drive early polarization.

**Figure 7.**
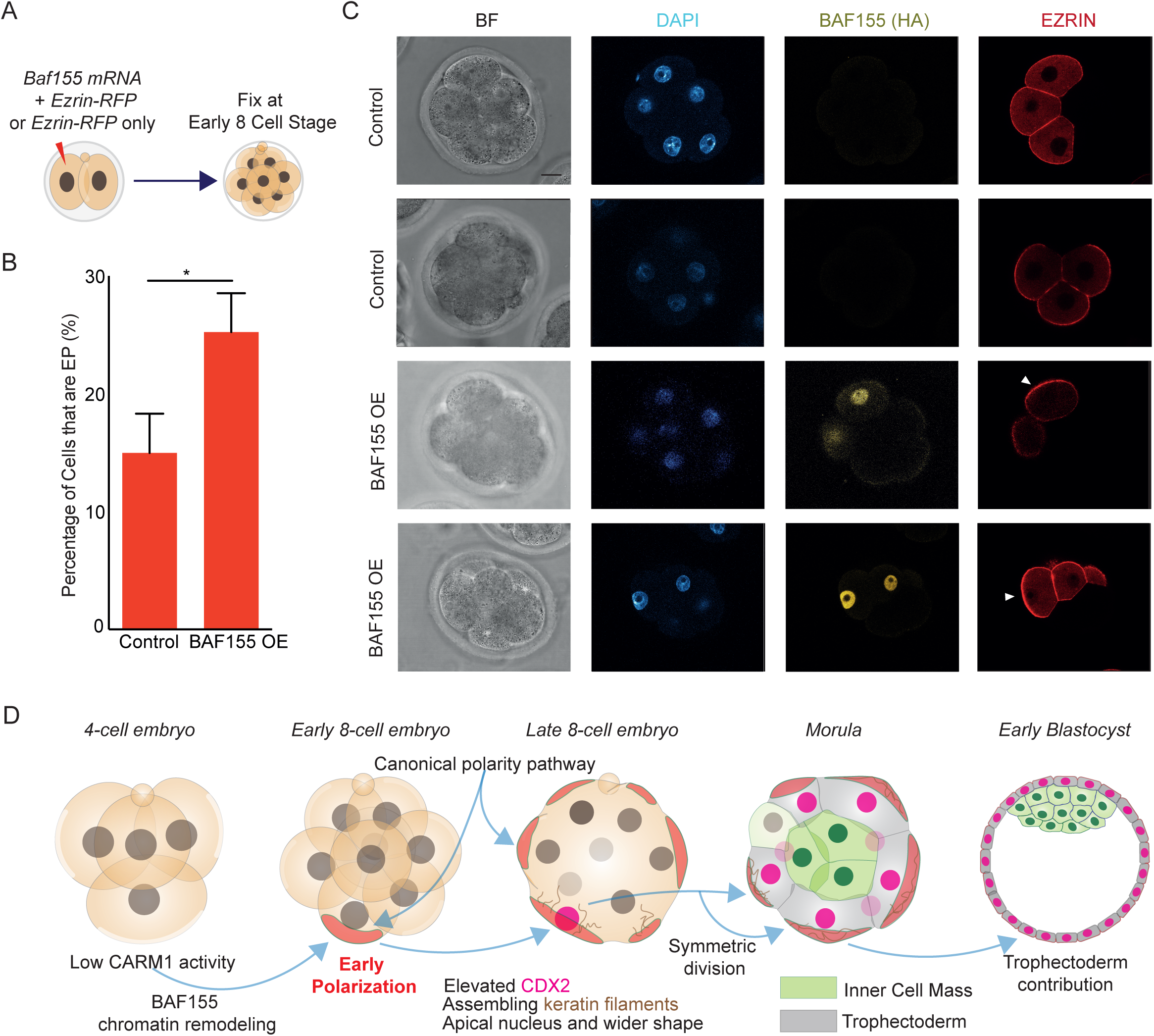
Frequency of early polarization is increased after BAF155 upregulation. A) Schematic indicating experimental design for overexpression (OE) of *Baf155* mRNA. B) BAF155 OE blastomeres have a significantly higher frequency of being early polarized than Control (*Ezrin*-RFP only) injected blastomeres at the early 8-cell stage, two-tailed z-test, *p < 0.05; N = 3 independent experiments, overall n = 35 BAF155 OE embryos with n = 140 cells (35 EP, 25.0%), n = 30 Control embryos with n = 120 cells (16 EP, 13.3%). Mean of individual experiments with S.E.M given on the graph. C) Immunostaining for DAPI, EZRIN and HA-tag (tagged to the BAF155 construct) at the early 8-cell stage. EZRIN-RFP present in injected blastomeres and an EP blastomere is indicated via an arrow. D) Schematic representing findings in this study integrating concepts from other studies. Blastomeres in the early embryo with low CARM1 activity (Torres-Padilla et al., 2007; Goolam et al., 2016)and elevated levels of its downstream target BAF155 (Panamarova et al., 2016) are biased towards polarizing early. Early polarization occurs through the canonical RHO A, TEAD4 and TFAP2C pathway (Zhu et al., 2017, 2020) and involves many of the same apical proteins, however these cells are distinguished by a number of factors which are consistent with its bias towards symmetric cell divisions and the trophectoderm lineage: early upregulation of Yap and activation of CDX2 (Jedrusik et al., 2008; Skamagki et al., 2013), an apical nucleus (Ajduk et al., 2014), wider geometry (Niwayama et al., 2019) and assembly of keratin filaments(Lim et al., 2020). Scale Bar = 15 µm.

Blastomeres with elevated BAF155 levels have been shown to form filaments of KERATIN-8 and KERATIN-18 at the late 8-cell stage, which helps stabilize the apical domain and promote TE specification. We stained early and late stage 8-cell embryos, as well as blastocysts, with KERATIN-18 (KRT18) and found that KRT18 positive cells emerged and increased in number from the late 8-cell stage onward (Figure 7-figure supplement 1A-B) (Lim et al., 2020).

Intriguingly, we found that approximately 7% of cells in the late 8-cell embryo form KRT18 filaments, which corresponds to the average proportion of early polarizing blastomeres. Finally, to test if early polarizing cells have precocious expression of *Krt18* mRNA, we performed single cell RNA sequencing on early and unpolarized blastomeres isolated from the early 8-cell stage (Figure 7-figure supplement 1C). We found that compared to unpolarized blastomeres, early polarizing blastomeres displayed a significant upregulation of *Krt18* mRNA alongside slight upregulation of *Baf155* mRNA (Figure 7-figure supplement 1D). These results suggest that low CARM1 activity and elevated BAF155 expression can enable blastomeres to polarize early.

## Discussion

All blastomeres of the mouse embryo become polarized at the late 8-cell stage. By studying the formation of the apical domain in detailed time-course studies, we found that blastomere polarization is asynchronous, with some blastomeres polarizing at the early 8-cell stage, before embryo compaction, while majority of blastomeres polarize after embryo compaction. These early polarizing blastomeres show distinct cellular behaviors compared to blastomeres that polarize late, including a closer nucleus-cortex distance, upregulation of CDX2 expression, a bias towards symmetric cell divisions and consequent differentiation into the TE.

It has been reported that blastomeres of the mouse embryo show differential activity of CARM1 at the 4-cell stage, which impacts their fate (Torres-Padilla et al., 2007; Goolam et al., 2016; Hupalowska et al., 2018). Here we show that reduced CARM1 activity and upregulation of its target BAF155 at the 4-cell stage promote early polarization, suggesting that the heterogeneous timing of embryo polarization is linked to earlier heterogeneities in the embryo. Previous work showed that the inhibition of CARM1 activity results in an increased number of symmetric cell divisions, as well as elevated CDX2 expression, which are consistent with the phenotypes present in early polarizing blastomeres (Parfitt and Zernicka-Goetz, 2010; Goolam et al., 2016). We show here that manipulating the levels of heterogeneously expressed cell fate regulators (CARM1, BAF155) alters the timing of polarization and emergence of the apical domain in the 8-cell embryo. These results suggest a link between asymmetries at the 4-cell stage and timing of blastomere polarization at the 8-cell stage, two mechanisms of the first cell fate decision that had been considered until now separate (Panamarova et al., 2016; Torres-Padilla et al., 2007; Lim et al., 2020).

What biases the early polarizing blastomeres towards symmetric cell divisions and consequently the TE fate? Cell shape is known to influence division orientation and lineage bias (Gray et al., 2004; Korotkevich et al., 2017; Niwayama et al., 2019), and the TE bias of early polarizing cells can be explained by an altered apical domain size, cell shape and increased symmetric cell divisions, as well as an early upregulation of TE determinants such as YAP and CDX2 and a smaller apical-nucleus distance, compared to cells that polarize late. Thus, our results allow us to link together multiple elements previously reported (Torres-Padilla et al., 2007; Goolam et al., 2016; Panamarova et al., 2016; Lim et al., 2020; Korotkevich et al., 2017; Niwayama et al., 2019; Ajduk et al., 2014; Jedrusik et al., 2008; Parfitt and Zernicka-Goetz, 2010) to explain the first cell fate decision (Figure 7D). It was known that 4-cell blastomeres have heterogeneous CARM1 activity and its downstream target BAF155 (Torres-Padilla et al., 2007; Panamarova et al., 2016; Lim et al., 2020; Piotrowska-Nitsche et al., 2005). Our results indicate that CARM1 and BAF155 can alter the timing of embryo polarization to influence subsequent cell fate: low CARM1 activity and high BAF155 levels promote early polarization, and early polarizing cells have altered nuclear position, cell geometry, CDX2 and keratin expression, that are each associated with symmetric distribution of the apical domain after its re-aggregation at the 16-cell stage, and subsequent bias towards the TE lineage.

The results we present here help to explain the TE fate bias of the early polarizing blastomeres, but they also help to explain the cellular mechanisms for the previously known role of CARM1 in promoting pluripotency and suppressing TE fate (Goolam et al., 2016; White et al., 2016). Given the regulative nature of mouse embryos, and the need for segregating the correct number of cells to each lineage (Morris et al., 2010), it is unsurprising that multiple mechanisms combine to influence cell fate. It is interesting to speculate that these findings in mouse embryos might have implications for other mammals, such as rabbits, cows and humans, which display substantial heterogeneity in the timing of blastomere polarization (Koyama et al., 1994; Zhu et al., 2021). Although the causes and consequences are unknown, it is possible that these differences are related to heterogeneities in early development and have effects on cell fate, respectively. Further insight into how polarization timing impacts the cell fates may have clinical implications as polarization is important for correct establishment of the blastocyst and human pregnancies often fail at the pre-implantation stage (Alarcon et al., 2010).

## Materials and Methods

### Availability

Commercially available materials are listed in the ‘Key Resources Table’ and their use is detailed in subsequent sections. All other materials are lab constructs, whose design and use has been detailed in the following sections. Upon reasonable request, these resources can be shared.

## Key Resources Table

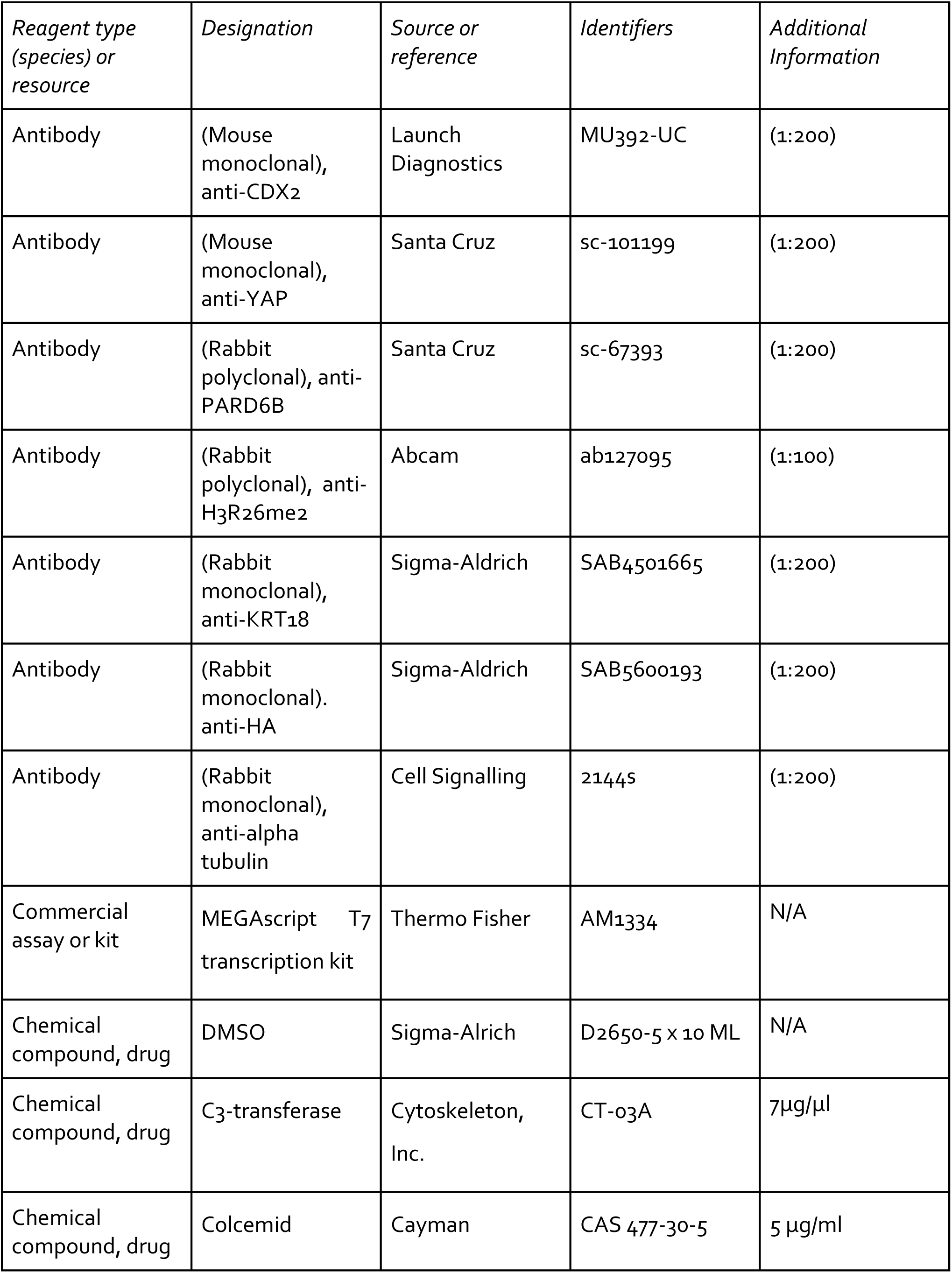
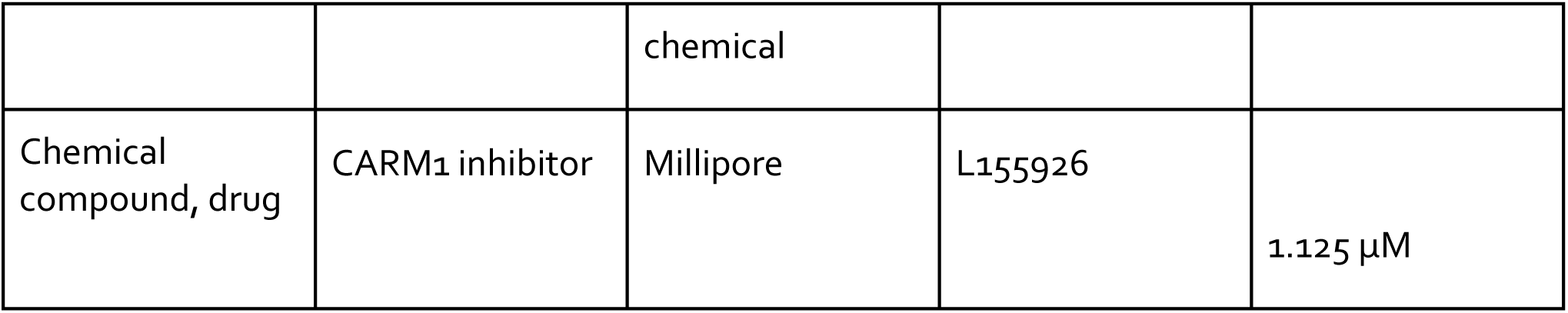

### Animals

The work presented here has been carried out under the regulations listed in the Animals (Scientific Procedures) Act 1986 [Amendment Regulations 2012] and further reviewed by the University of Cambridge Animal Welfare and Ethical Review Body. Embryos were collected from F1 females (C57BI6xCBA; Charles River strain 616) mated with F1 studs unless otherwise specified. Unless otherwise specified, 4-week old female mice were super-ovulated by injection of 7.5 IU of pregnant mares’ serum gonadotropin (Intervet, PMS) followed after 48 hours by human chorionic gonadotropin (Intervet, HCG), and mated with studs on the same day as HCG injection for subsequent embryo collection. When specified, natural mating of mice involves pairing a female with a stud overnight and subsequently collecting embryos post-mating.

### Embryo Culture and Treatment

Embryos were recovered in M2 medium (prepared in the lab or M7167; Sigma-Aldrich) at the required stage (2-cell stage, E1.5, unless specified) by tearing of the oviduct, and subsequently transferred to EmbryoMax® Advanced KSOM medium (MR-101-D-Aldrich) for long-term culture in 5.0% CO_2_ and 37 degrees Celsius, as described previously (Zhu et al., 2017). Individual variability in culture post-recovery was present as a result of the further experimental procedures required; it is specified for each experiment and described further in this section. Embryos remained in KSOM unless specified.

Disaggregation of 4-cell blastomeres in order to make 8-cell doublets and single cells for RNA sequencing was achieved as previously described (Johnson et al., 1986). Briefly, 4-cell stage embryos were washed in Acid Tyrode’s solution to remove the zona pellucida, before being separated by repeated pipetting in magnesium and calcium-free M2 with a flame-polished glass pipette.

C3-transferase was dissolved in distilled water and diluted in KSOM to 7μg/μl. Control treatment was KSOM with equivalent amount of vehicle (water) only. C3-transferase was applied to the embryo during the 4-8 cell stage division, before fixation and imaging.

Colcemid treatment has been previously used to depolymerize microtubules in the embryo (Sutherland and Calarco-Gillam, 1983). Reduced system cells currently in the 4-8 cell division stage were treated with colcemid in DMSO (5 μg/ml) for the duration of live imaging. Control treatment was DMSO only.

We inhibited the activity of CARM1 using a previously used drug (Cheng et al., 2011; Panamarova et al., 2016) at a concentration which was verified as influencing H3R26me2 activity whilst also allowing embryo survival (1.125 µM), dissolved in DMSO and diluted to its final concentration in KSOM. This was compared to a control condition in which we used the same volume of DMSO only instead.

### Preparation of Constructs

The following constructs were prepared for mouse embryo injection from primers and reagents previously used in our lab (Zhu et al., 2017; Parfitt and Zernicka-Goetz, 2010; Panamarova et al., 2016): Ezrin-RFP mRNA (Ezrin-RFP), *Tead4* dsRNA (*dsTead4*), *Tfap2c* dsRNA (*dsTfap2c*), *Carm1* E267Q mRNA (CARM1 E27Q), *Baf155* mRNA (*BAF155 OE*), *LifeAct-GFP* mRNA (*LifeAct*-GFP).

The first step in mRNA synthesis was cloning of the gene cDNA into the relevant pRN3P vector (pRN3P or pRN3P-RFP), as previously described (Zernicka-Goetz et al., 1997), after amplification of target genes from mouse, liver or kidney cDNA using previously specified primers. Subsequently, the pRN3P constructs were linearized using a restriction site on the plasmid downstream of the poly-A region, before in vitro transcription via the mMessage mMachine kit, as per the manufacturer’s instructions. mRNA was purified using lithium chloride precipitation.

For dsRNA, the E-RNAi website (Horn et al., 2010) was used to design constructs of 350-500 base pairs in length. Primers for each dsRNA (*dsTead 4* and *dsTfap 2c*) based on this design were acquired from Sigma-Aldrich and used to amplify target gene regions from a mouse, kidney, liver cDNA mixtures. Next, in vitro transcription was performed with the MEGAscript T7 transcription kit (Thermo Fisher, AM1334) as per the manufacturer’s instructions. dsRNAs were purified by lithium chloride precipitation.

### Microinjection

The injection of prepared constructs into mice was carried out as previously described (Zernicka-Goetz et al., 1997), on both blastomeres of the 2-cell stage embryo unless specified (in some cases, one blastomere was injected in order to have an uninjected half of the embryo as a developmental control). In summary, following recovery, embryos were placed in an M2 drop covered in mineral oil on a glass slide with a depression (that holds the drop). The slide was placed on a stage and held in place by holding apparatus. Embryos were injected with fluid from an injection needle via the Eppendorf Femtojet Microinjector. Negative capacitance was used to facilitate penetration through the membrane. *dsTfap 2c* and *dsTead4* were injected at a concentration of 1μg/μl. *Ezrin-RFP* was injected at a concentration of 400 ng/μl. *Carm1(E267Q),* and *Baf15*5 OE were injected at concentrations of 700 ng/μl.

### Immunofluorescence

Immunofluorescence was conducted in a 96-well round bottom plate.

Embryos were first fixed in 4% PFA for 20 minutes at room temperature. Subsequently, embryos were washed three times in PBST (0.1% Tween in PBS, phosphate buffered solution). The embryos were then permeabilised in 0.5% Triton X-100 in PBS for 20 min at room temperature, further washed in PBST three times, and transferred to blocking solution (3% bovine serum albumin) for 2.5 hours at 4 °C. Embryos were then incubated with primary antibodies (diluted in blocking solution, concentrations given individually below) at 4 °C overnight.

After the incubation, embryos were washed twice in PBST and incubated with secondary antibodies (diluted in blocking solution, concentrations given individually) for 2 hours at room temperature with foil covering. Embryos were then stained with DAPI (diluted in PBST, 1:1000) for 15 min, followed by two washes in PBST.

Primary antibodies: rabbit polyclonal anti-PARD 6b (Santa Cruz, sc-67393, 1:200); mouse monoclonal anti-CDX2 (Launch Diagnostics, MU392-UC (Biogenex), 1:200); mouse monoclonal anti-YAP (Santa Cruz, sc-101199, 1:200); rabbit monocolonal anti-KRT 18 (Sigma-Aldrich, SAB4501665, 1:200), rabbit monoclonal anti-alpha tubulin (Cell Signalling, 2144s, 1:200), rabbit monoclonal anti-HA (Sigma-Aldrich, SAB5600193, 1:200), rabbit polyclonal anti-H3R26me2 (Abcam, ab127095, 1:100).

Secondary antibodies: Alexa Fluor 488 Donkey anti-Mouse, (A-21202, ThermoFisher Scientific); Alexa Fluor 568 Donkey anti-Mouse (A-10037, ThermoFisher Scientific); Alexa Fluor 647 Donkey anti-Mouse (A-31571, ThermoFisher Scientific); Alexa Fluor 568 Donkey anti-Rabbit (A-10042, ThermoFisher Scientific); Alexa Fluor 647 Donkey anti-Rabbit (A-31573, ThermoFisher Scientific).

F-ACTIN was stained by secondary antibody Alexa Fluor Phalloidin 488 (ThermoFisher Scientific, A-12379) alone, and nuclear staining was given by DAPI (Life Technologies, D3571).

### Hybridization Chain Reaction (HCR)

Sequential HCR immunohistochemistry and HCR RNA fluorescent in situ hybridisation was carried out on pre-implantation embryos in a 96-well plate as per the following protocol. https://www.molecularinstruments.com/hcr-ihc-hcr-rnafish-protocols.

Briefly: after fixation in PFA, the embryos were washed and permeabilised as for immunofluorescence. The embryos were placed in blocking buffer (antibody buffer) for 4 hours at 4 °C before being kept at the same temperature overnight in primary antibody in antibody buffer at listed concentration for immunofluorescence.

Embryos were then washed in PBST and SSCT (0.1% Tween in 5 X SSC, sodium chloride sodium citrate) before post-fixation in PFA, and then placed for pre-hybridization in pre-warmed probe hybridization buffer for 30 minutes at 37 °C before being kept overnight in a 16nM probe solution (*Cdx2*, *Nanog* or *Tead4*) in hybridization buffer at the same temperature. The next day, the probes were washed off with pre-warmed wash buffer and SSCT before being pre-amplified in amplification buffer at room temperature. 15pmol of each amplification hairpin (h1 and h2 corresponding to each channel and probe) were snap cooled in separate tubes by heating to 95 °C for 90 seconds before cooling in a dark room at room temperature for 30 minutes, before the embryos were placed in amplification solution of 60 nM hairpins diluted in amplification buffer overnight at room temperature. The embryos were then washed in SSCT before imaging.

### Imaging

Live imaging was performed as previously described (Zhu et al., 2020). Briefly, time-lapse recordings of embryos were carried out using a spinning disk or a Leica-SP5 scanning confocal. Live-imaging time lapse frames were acquired every 20 minutes unless specified, with conditions of 5.0% CO_2_ and 37 degrees Celsius. For fixed samples, embryos were imaged on a Leica-SP5 scanning confocal microscope. Images were taken under a Leica 1.4 NA 63X oil (HC PL APO) objective. Images were subsequently analysed on Fiji software (Schindelin et al. 2012) as specified in following sections.

For live imaging our embryos, we used glass-bottom 35 mm dishes. We then fixed a small cut square of nylon mesh (5mm to 1cm width and height) onto this plate in the centre using silicon which was used as a grid (diameter of approximately 150 micrometres) for deposition of embryos. After drying of the silicon (overnight) and washing with water, the grid was overlaid with a drop of 100 microlitres of KSOM and then covered with mineral oil until this KSOM drop was submerged. After incubation under conditions for live imaging, single embryos were deposited in each ‘well’ of the grid before being placed in the microscope, which was equilibrated at the correct temperature and CO2.

### Defining Polarization and Compaction

We define the apical domain based on three criteria: length, intensity and depth:

1. The length of the domain must be between 33% and 80% of the total contact-free length of the cell.
2. The mean intensity of this selected length must be more than 1.5X the mean intensity of the remaining contact-free length of the cell.
3. The apical domain must be visible across at least 3 µm of depth (z-axis).

For all live imaging experiments in which the timing was precisely measured, we evaluated the timing of polarization from the end of the 3^rd^ cleavage of each cell (two distinct cells visible) in order to determine the time of polarization of each individual cell.

For embryo-level experiments using fixed samples, we fixed embryos within an hour after division to the 8-cell stage, which we could accomplish by briefly checking the embryos at regular intervals (30 minutes). Embryos with significantly different developmental timing from the group (not at the 8-cell stage or compacted beyond 120 degrees inter-blastomere angle) were excluded.

Compaction is a gradual process which beings at the early-mid 8-cell stage, and where the inter-blastomere angle – the angle formed between the outer edge of two cells at the point of contact – shifts from being more acute (60 degrees) to being more obtuse (180 degrees) over time. An embryo can be considered compacted when all the inter-blastomeres angle reach a minimum of 120 degrees (half-way).

For live imaging experiments, we defined early polarizing (EP) cells as a cell that forms the apical domain (1) within the first hour after its 3rd cleavage and (2) before the inter-blastomere angle between the cell and its neighbours reached 120 degrees, in the process of blastomere compaction. Polarized cells not fulfilling these criteria were defined as ‘late polarizing’ (LP). For fixed embryos, we fixed immediately after the last blastomere took the 3rd cleavage division that generates the 8-cell embryo. When using this approach, we defined an EP cell as a cell that forms an apical domain (1) within the first hour after the last 3rd cleavage in this embryo and (2) before the inter-blastomere angle between the cell and its neighbours reached 120 degrees. Polarized cells not fulfilling these criteria were defined as ‘late polarizing’ (LP).

In both live and fixed embryos, if the blastomere polarizes after the inter-blastomere angle between the blastomere and its neighbours reaches 120 degrees, it is considered an LP, and embryos with at least one EP cell were deemed EP embryos.

### Data Processing and Statistics

Randomisation: Females and males were randomly chosen from a pool of animals. Embryos from different breeding pairs were recovered and mixed before they were randomly allocated to the different experimental groups.

Sample size: The size of the sample per condition in an experiment was determined empirically having as a reference other mouse embryo studies. In all cases unless otherwise specified, more than 15 embryos per condition were used.

Exclusion criteria: Embryos that showed apparent developmental delay or cell death were discarded.

Blindness: The investigators were not blinded to allocation during experiments and outcome assessment.

This research features both qualitative and quantitative data. For qualitative data, methods of analysis have been carefully described in text and figure captions to be as objective as possible. For quantitative data, a summary of statistical methods used in analysis are indicated for every experiment in the corresponding figure legends.

In general, quantitative data was analysed as follows: the fit of the data to a normal distribution was analysed with D’Agostino’s K-squared test. If data fit a normal distribution, then for comparison of two or multiple samples, an unpaired or paired two-tailed student’s *t* test (two experimental groups) or a one-way ANOVA test (more than two experimental groups) was used to analyse statistical significance, depending on the fit of the data to the assumptions of each test. Differences in variances were taken into account by performing a Welch’s correction where appropriate. For data that did not display a normal distribution, a Mann–Whitney *U*-test (for two experimental groups) or a Kruskal–Wallis test with a Dunn’s multiple comparison test (for more than two experimental groups) was used to test statistical significance. To determine the influence of different groups in multiple variants, a two-way ANOVA was performed. Statistical analyses were performed using the Graphpad Prism software (http://www.graphpad.com).

The probability distribution of EP cells according to cell autonomy is described in relevant text and figures.

Cortical signal enrichment of EZRIN was calculated by first measuring the signal intensity of the cortical line on Fiji (Icortex), and then a standard portion of the cytoplasm as control (Icytoplasm), before applying the formula: EZRIN cortical enrichment = Icortex−Icytoplasm)/Icytoplasm.

The IEA between adjacement blastomeres was calculated as previously described (Zhu et al., 2017; Fierro-Gonzalez et al., 2013). PARD6 and F-ACTIN enrichment was calculated by measuring cortical signal intensity over a 35 µm stretch of membrane in an EP and LP cell in the same z-axis, plotted and smoothened factor 20 with a 2^nd^ dimension polynomial on Graphpad Prism (http://www.graphpad.com).

Apical domain size was calculated as described in the corresponding figure. The angle subtended by this domain at the edge of the cell was then taken as the apical domain size in degrees, with the radius equal to the arc (see figure). This was done on the z plane of the embryo for which the nucleus was largest (approximately mid-plane).

Nucleus-apex distance was measured as the distance from the line made by the top of the nucleus to the approximately parallel line made by the centre of the outside surface of the cell. This was done on the z plane of the embryo for which the nucleus was largest (approximately mid-plane).

From time-lapse movies of development, the moment used for analyses of the cell size was the time point before the first blastomere began to divide at the 8-16 cell stage (if the blastomeres divided in adjacent frames) or the time point before the blastomere divided itself (if the cells divided at different times), and the plane used for each cell was that in which the nucleus was biggest (approximately mid-plane). The apex angle is the angle subtended at the centre when the edges are joined via converging lines, and the length ratio is the ratio of the average side length to the average length of the converging lines subtended at the centre. All geometries were analysed on Fiji.

Nuclear: cytoplasmic ratio was calculated by circling the nucleus and measuring nuclear signal intensity of the focal blastomere (INTnucleus), and then measuring intensity of a region of cytoplasm of the same size (INTcytoplasm), before performing the calculation: INTnucleus/INTcytoplasm. This was done on the z plane of the embryo for which the nucleus was largest (approximately mid-plane). Nuclear:cytoplasmic ratio in other instances was calculated in the same way.

Average (EP) intensity level: average (LP) intensity level ratio for each embryo was calculated as follows: First, nuclear intensity was measured by circling the nucleus of the z plane for which the nucleus was largest (approximately mid-plane). Then, nuclear DAPI intensity was measured for the same area. The nuclear signal: nuclear DAPI ratio was found for each blastomere of the embryo. Subsequently, the result for EP blastomeres was averaged, and likewise for the LP blastomeres, before finding the Average (EP) level: average (LP) level ratio for the particular embryo.

To assess CDX2-GFP signal in our doublet “8-cell stage” embryos, we measured nuclear signal in the mid-plane where both nuclei were visible over time and normalised this to the background (cytoplasmic) signal. *Cdx2* transcript counts were done manually in ImageJ by counting each individual punctum in cells over the same z-slices and Deng et al.(Deng et al., 2014) used as a reference.

3D analysis was conducted with the use of the Fiji Mesh Deformation software as described in the following page: https://imagej.net/plugins/deforming-mesh-3d. At each 8-cell stage time-point, a mesh was created for each cell. The information was then processed in Python in order to extract the relevant features including the cellular aspect ratio. The aspect ratio for each cell was normalized to the embryo average for the final quantification.

Other calculations were made from observations, as discussed in the text attributed to each figure and experiment. All visible blastomeres in an embryo were counted unless otherwise specified in the text. All scale bars represent 15 microns unless otherwise specified.

## Data availability

Source data is available from the corresponding author upon reasonable request.

## Acknowledgments

We would like to acknowledge all members of the Zernicka-Goetz lab and Angela Andersen from Life Science Editors for their support. We also thank our funding sources: Wellcome Trust (098287/Z/12/Z) (MZG), Leverhulme Trust (RPG-2018-085) (MZG), NIH R01 HD100456-01A1 (MZG), Medical Research Council (AL), Cambridge Vice Chancellor’s Award Fund (AL). There are no competing interests.

---

**Figure 1-figure supplement 1.**
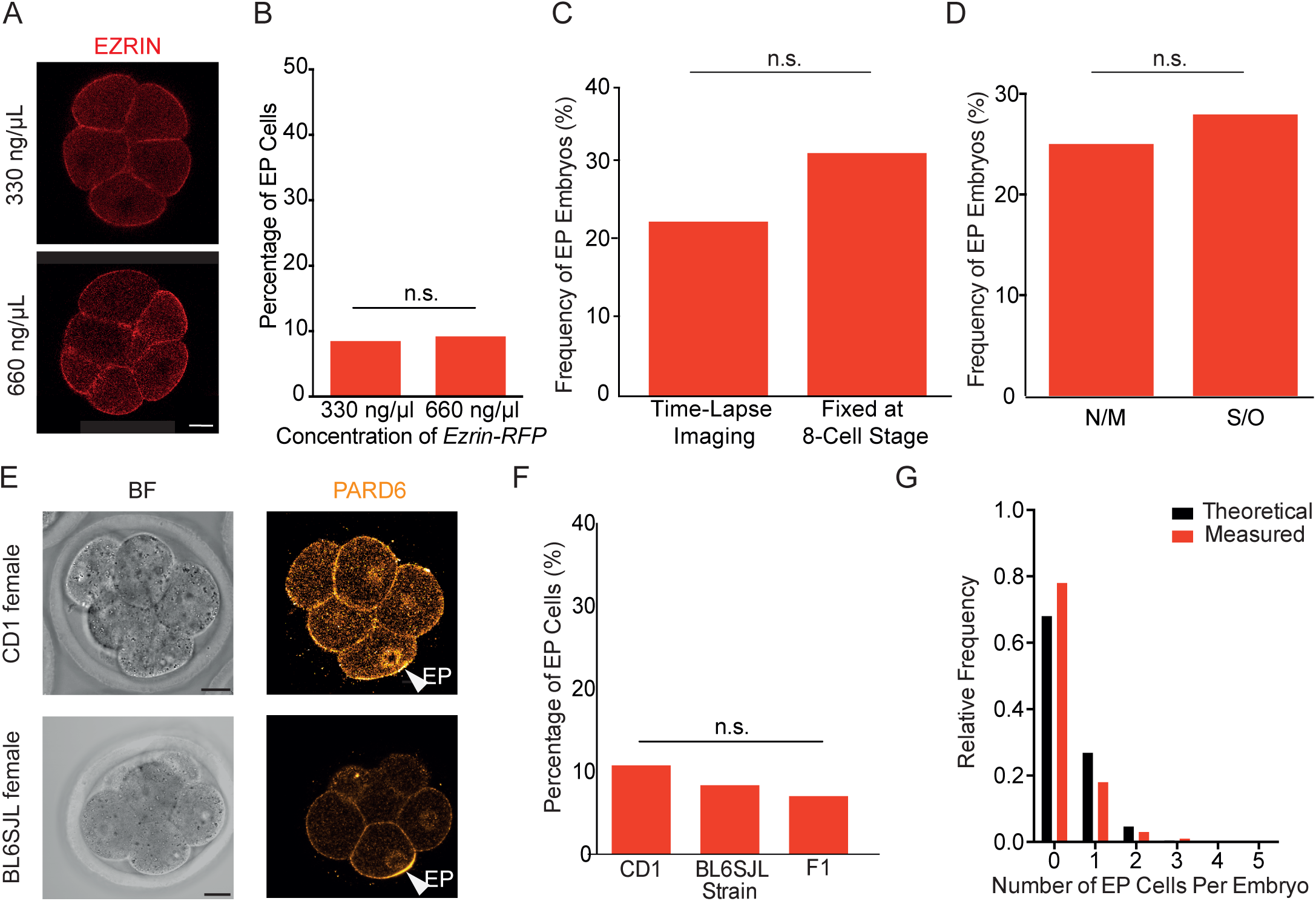
Early polarization is not dependent on imaging or mating type, or embryo effects. A) Representative images showing endogenous EZRIN-RFP signal in embryos injected with 330 ng/μl or 660 ng/μl. B) The concentration of injected mRNA does not affect the frequency of early polarizing (EP) cells, n = 82 blastomeres counted for 330 ng/μl from 11 embryos (7 EP cells), n = 75 blastomeres counted for 660 ng ng/μl from 10 embryos (7 EP cells), two-tailed z-proportion test, n.s.is non-significant, p > 0.05, N = 3 independent experiments. C) The frequency of early polarizing embryos does not differ significantly between time-lapse imaging and fixed embryos, two-tailed z-proportion test, n.s.is non-significant, p > 0.05, n= 87 embryos analyzed in time-lapse videos (19 EP), n = 32 fixed samples (10 EP). An EP embryo is one that contains at least one EP cell, as defined in Materials and Methods. Time-lapse imaging was conducted with microinjection of Ezrin-RFP into blastomeres, at the 2-cell stage, and embryos were imaged from the 8-cell stage, with blastomeres polarising in the first hour being counted as EP. For fixed embryos, Pard6 was used as the marker and embryos were fixed one hour after division to the 8-cell stage. Please see Materials and Methods for further details. D) Frequency of early polarizing embryos does not differ significantly between naturally mated and super-ovulated mice, n = 87 super-ovulated embryos (19 EP), n = 16 naturally-mated embryos (4 EP). two-tailed z-proportion test, n.s.is non-significant, p > 0.05. E) Representative immunostaining images (using PARD6 as a polarity marker) of embryos from super-ovulated CD1 and BL6SJL females at the 8-cell stage. F) Quantification of percentage of EP cells in different mouse strains, n = 56 CD1 blastomeres (7 embryos), 6 EP cells (10.7%); n = 120 BL6SJL blastomeres (15 embryos), 10 EP cells (8.3%) with comparison to previously reported quantification in F1 mice (7.0% EP cells). There is no significant difference in percentage of EP cells in any strain from all pairwise comparisons, two-tailed z-proportion tests, n.s.is non-significant, p > 0.05. G) Theoretical number of EP cells per embryo assuming independent cell autonomy, calculated by 8Cn X P^n^ X (1-P)^8-n^ where p is the probability of an EP cell and n is the number of EP cells in the embryo, compared with measured values. Scale Bar = 15 µm.

**Figure 2-figure supplement 1.**
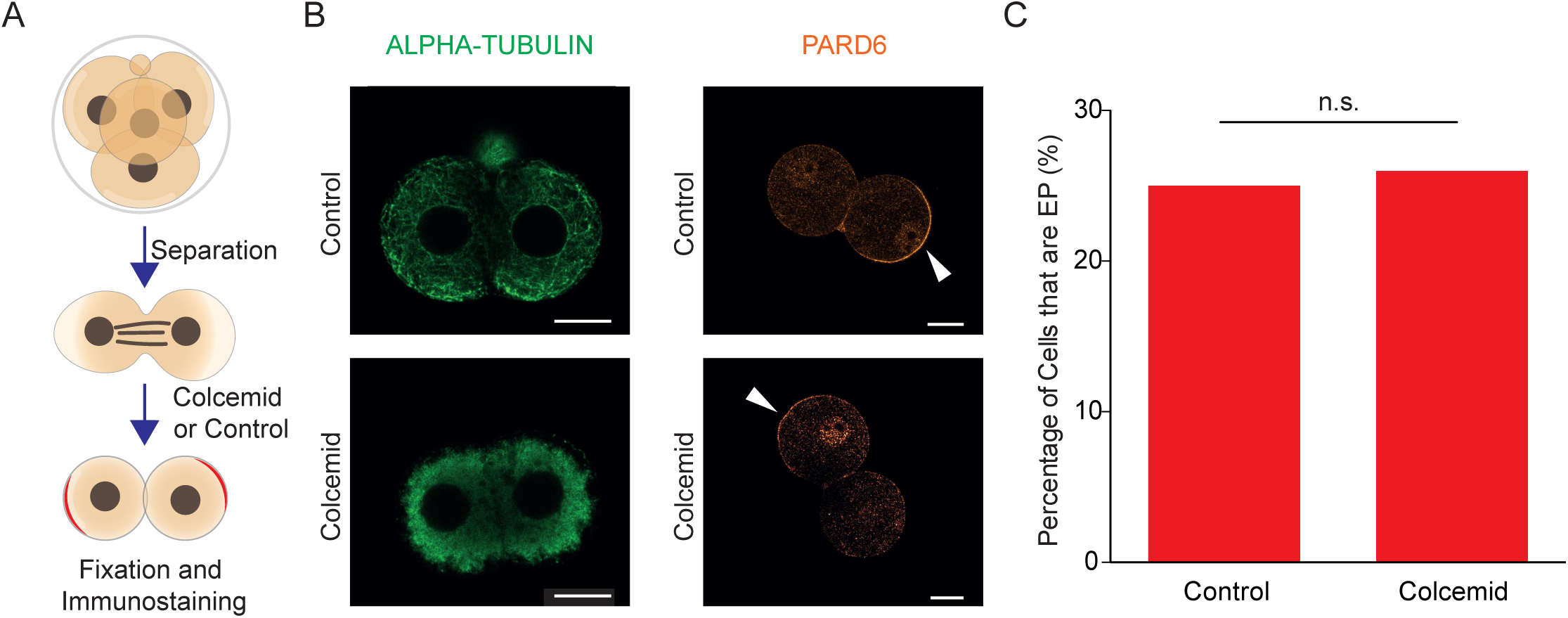
Microtubule depolymerization does not affect the frequency of early polarization. A) Schematic of disaggregating 4-cell stage embryos into individual blastomeres before treatment with colcemid or DMSO. B) ALPHA-TUBULIN and PARD6 immunostaining of early 8-cell doublets in control and colcemid-treated conditions. C) No significant difference between percentage of EP cells in control and colcemid-treated early 8-cell doublets, two-tailed t-test, p > 0.05, n = 49 colcemid-treated cells, n = 51 control-treated cells. Scale Bar = 15 µm.

**Figure 2-figure supplement 2.**
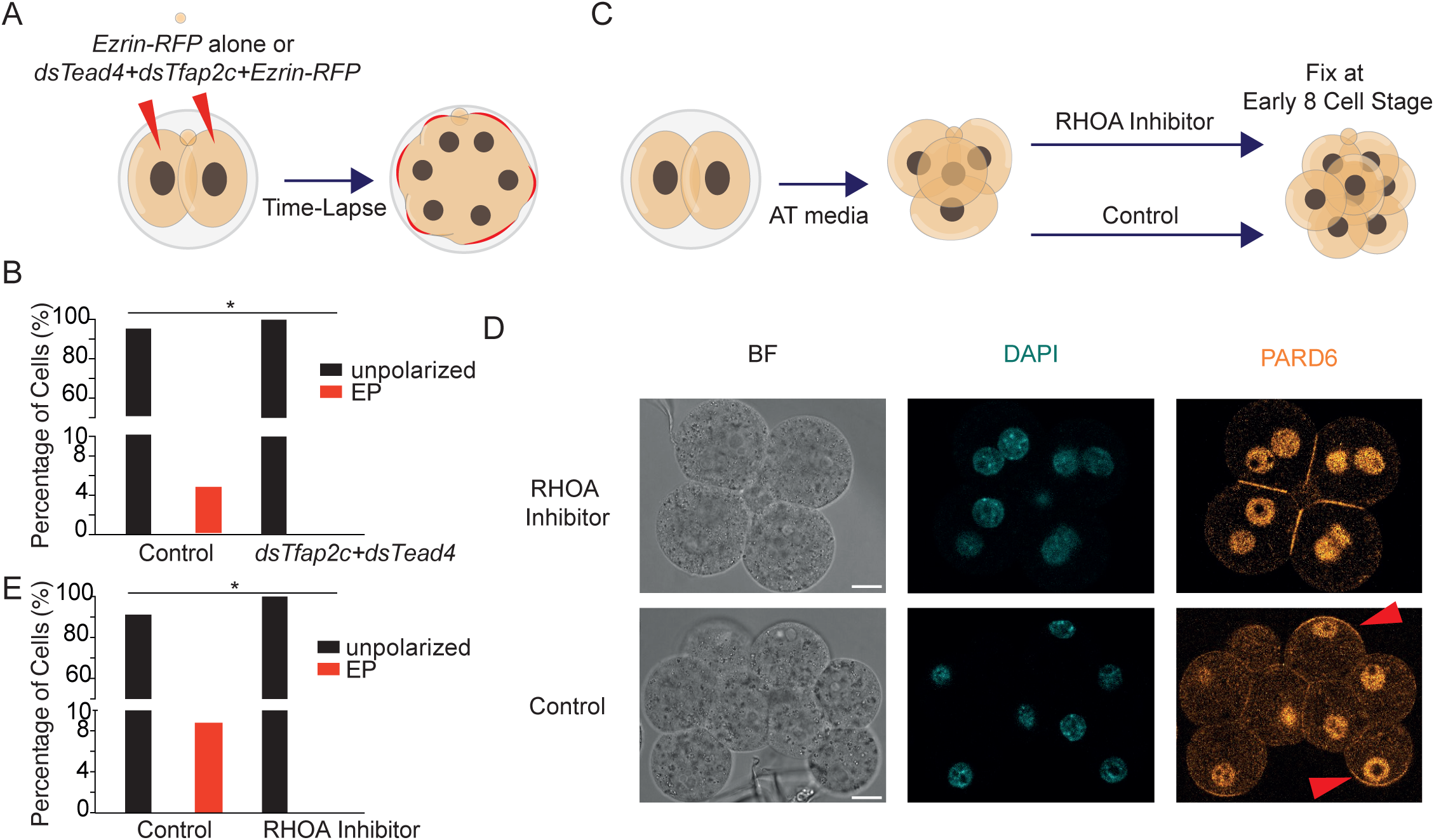
Early polarization requires TEAD4, TFAP2C and RHOA. A) Schematic indicating experimental design to deplete *Tfap2c* and *Tead4* simultaneously. B) Depletion of *Tfap2c* and *Tead4* simultaneously by dsRNA injection causes elimination of EP cells, N = 2 independent experiments, n = 17 dsTead4+dsTfap2c embryos with n = 134 cells (0 EP), n = 22 Control embryos with n = 170 cells (8 EP), overall percentages given on graph, two-tailed z-test, *p < 0.05. C) Schematic representing treatment of embryos with C3 Transferase to inhibit RhoA, or with a control treatment. D) Immunostaining images showing RHO A inhibited embryos versus control embryos stained for DAPI and PARD6, indicating binucleation of RHO A embryos and arrows pointing to EP cells. E) Inhibition of RHO A causes elimination of EP cells, N = 3 independent experiments, n = 8 RhoAi embryos with n = 32 binucleated cells (0 EP), n = 16 Control embryos with n = 125 cells (11 EP), overall percentages given on graph, each binucleated cell counted as two cells for statistical analysis, two-tailed z-test, *p < 0.05. Scale Bar = 15 µm.

**Figure 3-figure supplement 1.**
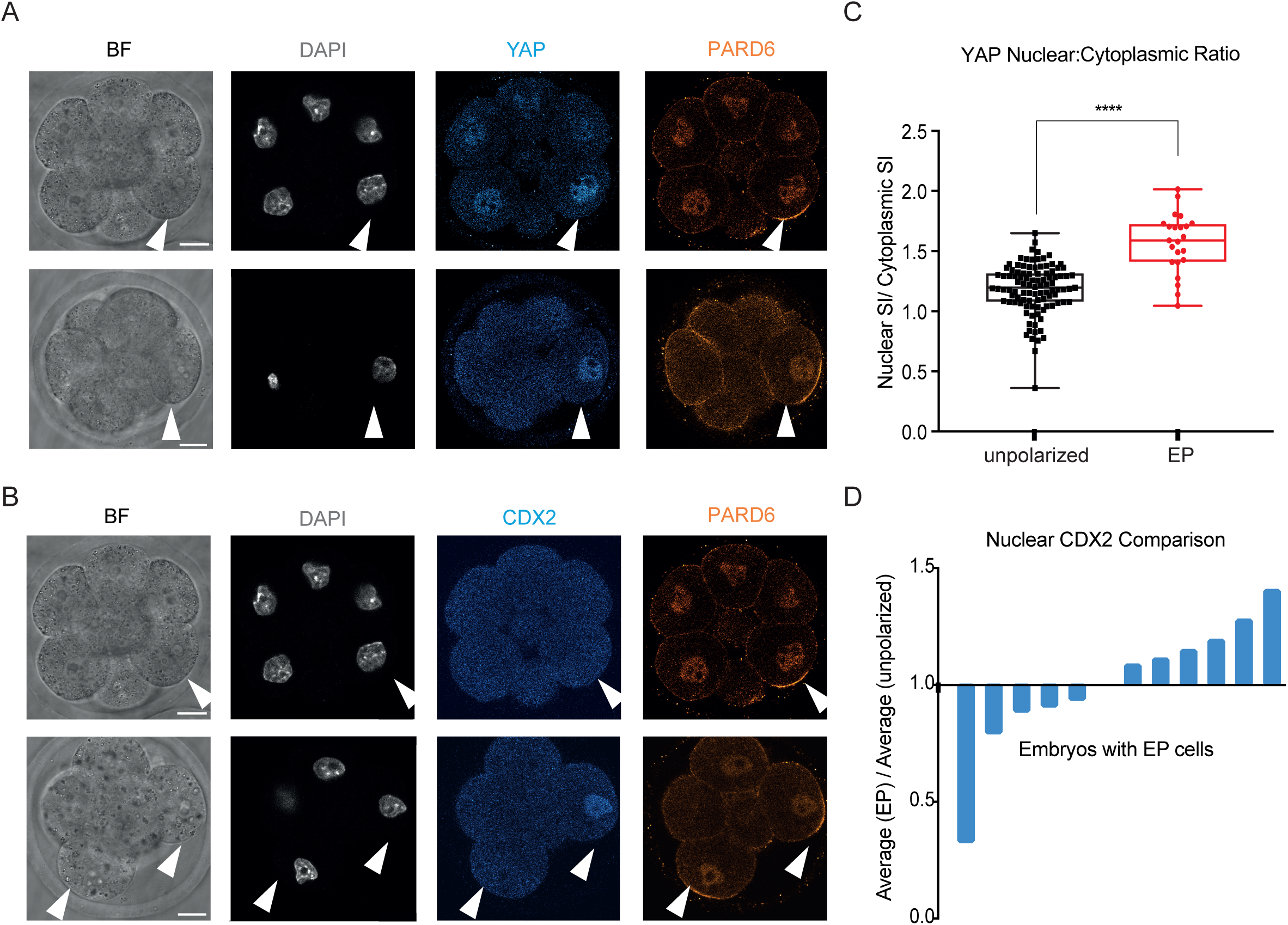
Nuclear YAP, but not nuclear CDX2, is higher in EP versus unpolarized blastomeres of uncompacted embryos. A) Immunostaining of uncompacted, early 8-cell stage embryos for YAP and PARD6. EP cells are indicated with an arrowhead and have apical PARD6 accumulation. B) Immunostaining of uncompacted, early 8-cell stage embryos for CDX2 and PARD6. EP cells are indicated with an arrowhead and have apical PARD6 accumulation. C) The nuclear:cytoplasmic ratio of YAP in EP and unpolarized blastomeres; two-tailed t-test, ****p < 0.0001; N = 4 independent experiments, n = 16 embryos, with n = 124 cells (23 EP). D) Ratio of average nuclear CDX2 in EP versus unpolarized blastomeres across different embryos. n = 12 EP embryos. Scale Bar = 15 µm.

**Figure 6-figure supplement 1.**
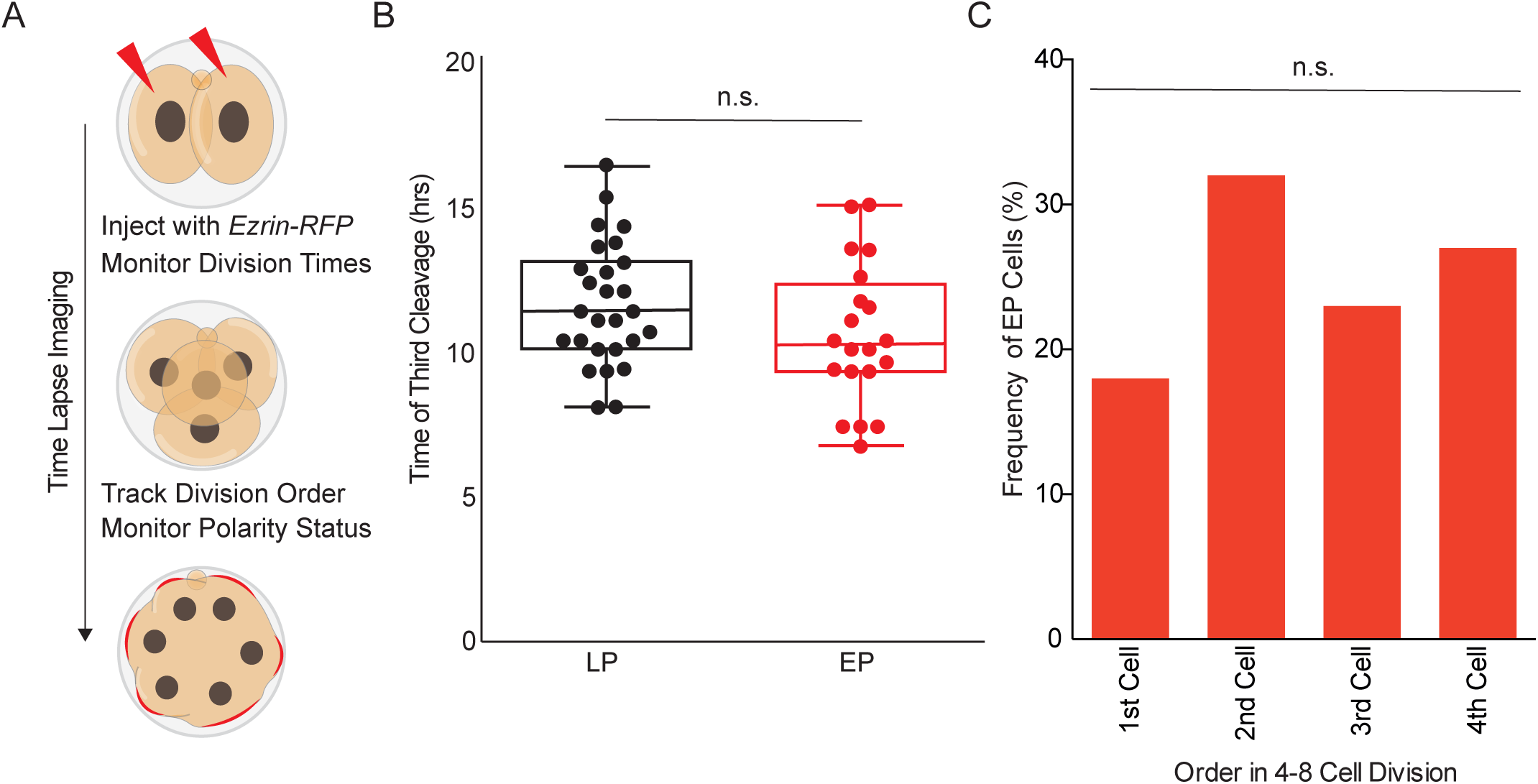
Early polarization is not dependent on division timing or order. A) schematic showing experimental design to measure division and polarization timing. B) No significant difference between EP and LP populations in timing of third cleavage, n = 47 cells assessed, two-sample t-test, n.s. – not significant p>0.05. C) Order of division is not correlated with frequency of EP cells, n = 22 cells assessed, histogram correlation computed, n.s. – not significant, p>0.05.

**Figure 6-figure supplement 2.**
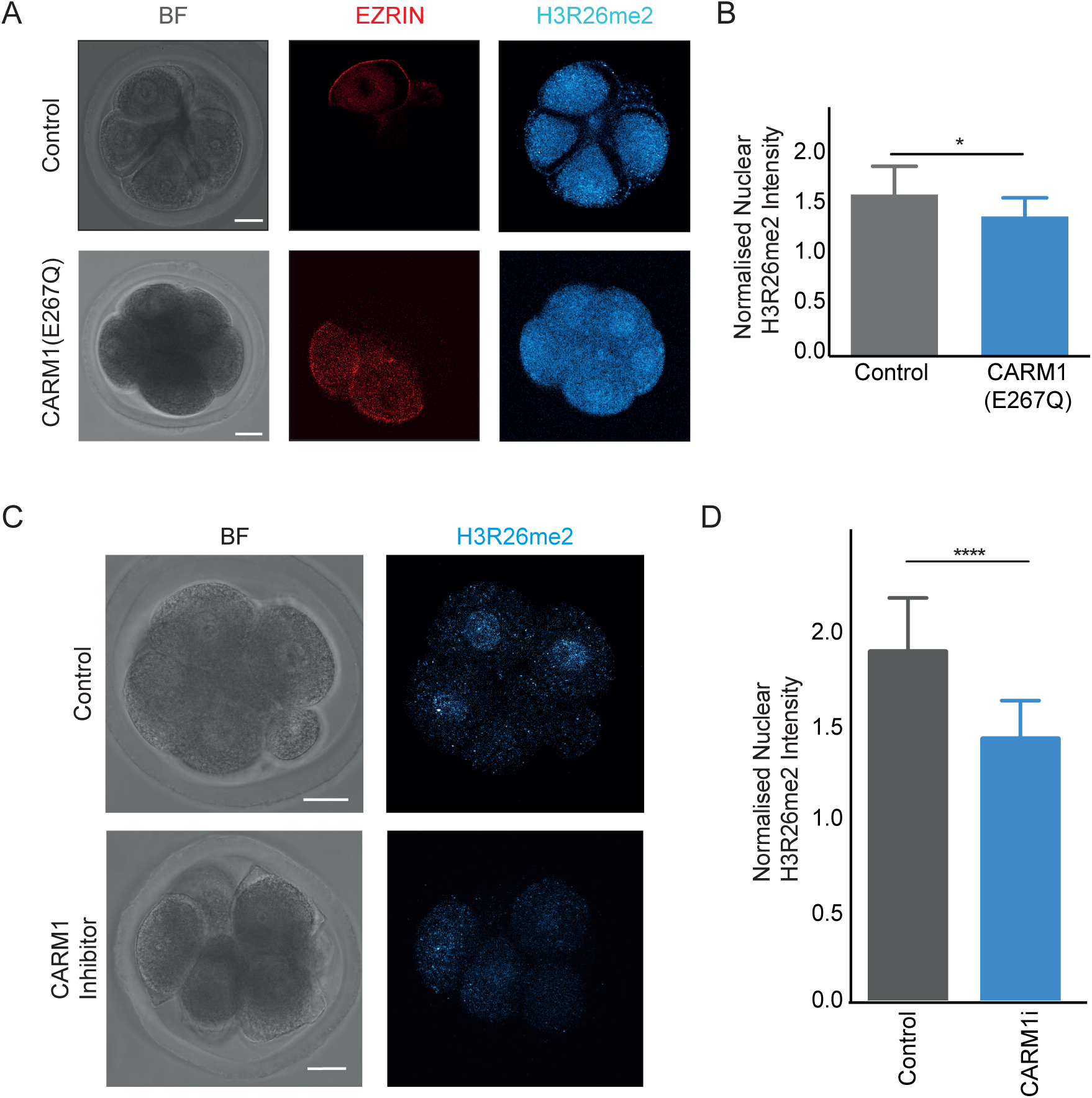
CARM1 (E267Q) and CARM1 inhibitor verification using H3R26me2 staining. A) Immunostaining for H3R26me2 (double methylation of arginine 26 on histone H3) in early 8-cell embryos injected with CARM1(E267Q). B) Quantification of nuclear H3R26me2 (normalized to background cytoplasm) shows significantly lower H3R26me2 nuclear intensity in CARM1(E267Q) blastomeres than Control blastomeres, two-sample t-test, **p<0.05, n = 10 Control blastomeres from 3 embryos and n = 15 CARM1(E267Q) blastomeres from 4 embryos. C) Immunostaining for H3R26me2 (double methylation of arginine 26 on histone H3) in early 8-cell embryos treated with CARM1 inhibitor or just vehicle. D) Quantification of nuclear H3R26me2 (normalized to background cytoplasm) between inhibitor and control treatments for 8 cell stage cells, showing a significantly higher concentration for the control treatment compared with CARM1 inhibitor, two-tailed t test, ****p < 0.0001, n = 3 inhibitor treated embryos (22 cells analyzed), n = 2 control treated embryos (14 cells analyzed). All graphs shown with standard error of the mean. Scale Bar = 15 µm.

**Figure 7-figure supplement 1.**
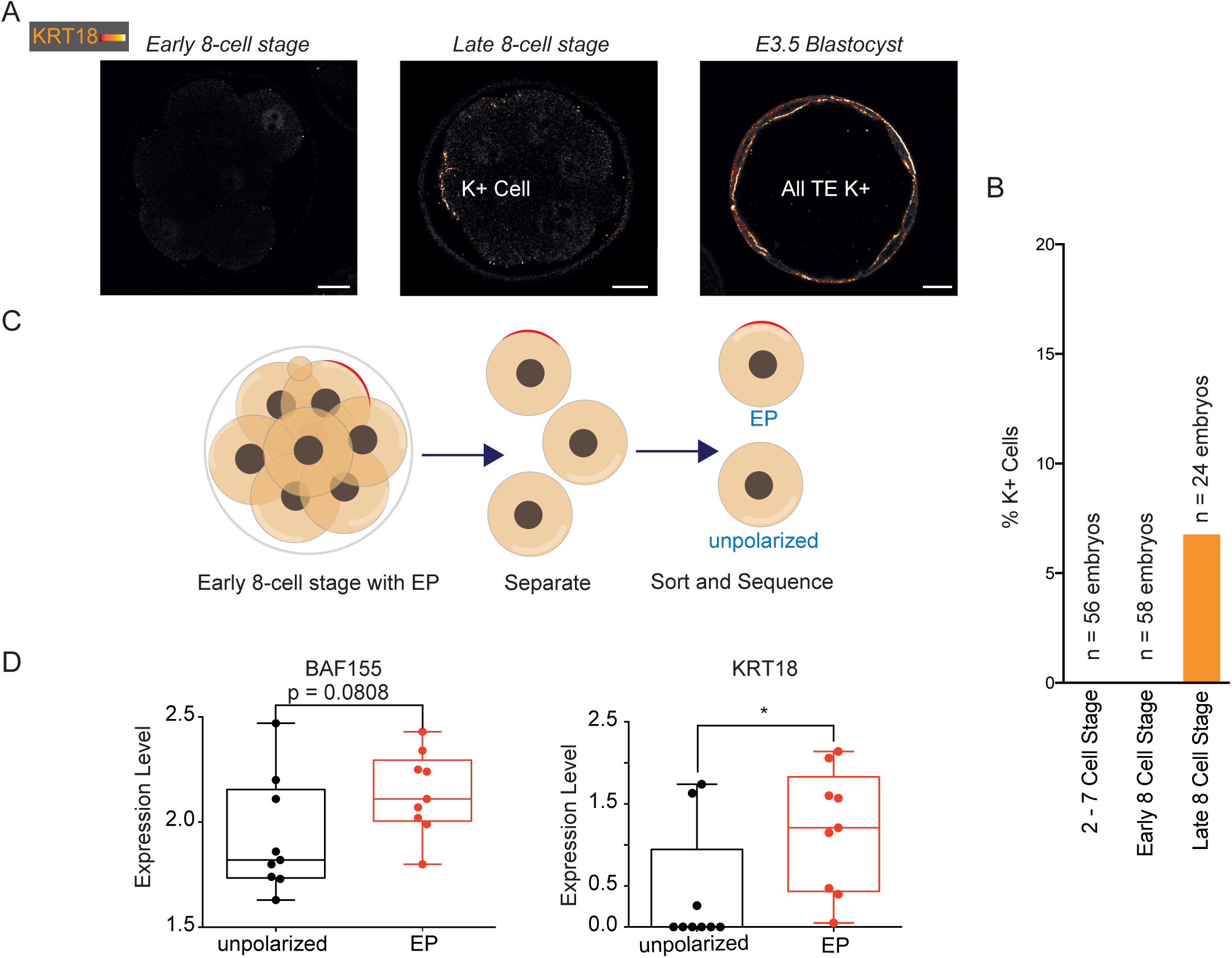
*Krt18* mRNA is upregulated in early polarized versus unpolarized blastomeres. A) Representative z-slice of immunostaining of early 8-cell stage, late 8-cell stage and E3.5 embryos for KRT18. B) Graph showing the frequency of KRT18-positive cells at different stages during early pre-implantation murine development. We found that 6.77% of cells at the late 8-cell stage show keratin accumulation. C) Schematic of workflow: early 8-cell embryos were disaggregated into individual EP and unpolarized blastomeres before single cell RNA sequencing. Polarity status was determined by using live imaging of EZRIN-RFP. 10 EP and 10 unpolarized cells sequenced D) Expression levels of *Baf155* and *Krt18* determined by single-cell sequencing, two-tailed t-test, p = 0.0808 for *Baf155* and p = 0.0394, *<0.05 for *Krt18*. Scale Bar = 15 µm.

## References

Ajduk, A., S. Biswas Shivhare, and M. Zernicka-Goetz. 2014. The basal position of nuclei is one pre-requisite for asymmetric cell divisions in the early mouse embryo. Dev Biol. 392:133–140. doi:10.1016/j.ydbio.2014.05.009.

Alarcon, V.B. 2010. Cell Polarity Regulator PARD6B Is Essential for Trophectoderm Formation in the Preimplantation Mouse Embryo1. Biol Reprod. 83:347–358. doi:10.1095/biolreprod.110.084400.

Anani, S., S. Bhat, N. Honma-Yamanaka, D. Krawchuk, and Y. Yamanaka. 2014. Initiation of Hippo signaling is linked to polarity rather than to cell position in the pre-implantation mouse embryo. Development (Cambridge). 141:2813–2824. doi:10.1242/dev.107276.

Bischoff, M., D.E. Parfitt, and M. Zernicka-Goetz. 2008. Formation of the embryonic-abembryonic axis of the mouse blastocyst: Relationships between orientation of early cleavage divisions and pattern of symmetric/asymmetric divisions. Development. 135:953–962. doi:10.1242/dev.014316.

Cheng, D., S. Valente, S. Castellano, G. Sbardella, R. Di Santo, R. Costi, M.T. Bedford, and A. Mai. 2011. Novel 3,5-Bis(bromohydroxybenzylidene)piperidin-4-ones as Coactivator-Associated Arginine Methyltransferase 1 Inhibitors: Enzyme Selectivity and Cellular Activity. J Med Chem. 54:4928– 4932. doi:10.1021/jm200453n.

Deng, Q., D. Ramsköld, B. Reinius, and R. Sandberg. 2014. Single-Cell RNA-Seq Reveals Dynamic, Random Monoallelic Gene Expression in Mammalian Cells. Science (1979). 343:193–196. doi:10.1126/science.1245316.

Ducibella, T., and E. Anderson. 1975. Cell Shape and Membrane Changes in the Eight-Cell Mouse Embryo: Prerequisites for Morphogenesis of the Blastocyst’. 47. 45–58 pp.

Dujardin, D.L., and R.B. Vallee. 2002. Dynein at the cortex. Curr Opin Cell Biol. 14:44–49. doi:10.1016/S0955-0674(01)00292-7.

Goolam, M., A. Scialdone, S.J.L. Graham, I.C. MacAulay, A. Jedrusik, A. Hupalowska, T. Voet, J.C. Marioni, and M. Zernicka-Goetz. 2016. Heterogeneity in Oct4 and Sox2 Targets Biases Cell Fate in 4-Cell Mouse Embryos. Cell. 165:61–74. doi:10.1016/j.cell.2016.01.047.

Gray, D., B. Plusa, K. Piotrowska, J. Na, B. Tom, D.M. Glover, and M. Zernicka-Goetz. 2004. First Cleavage of the Mouse Embryo Responds to Change in Egg Shape at Fertilization. Current Biology. 14:397–405. doi:10.1016/j.cub.2004.02.031.

Horn, T., T. Sandmann, and M. Boutros. 2010. Design and evaluation of genome-wide libraries for RNA interference screens. Genome Biol. 11:R61. doi:10.1186/gb-2010-11-6-r61.

Hupalowska, A., A. Jedrusik, M. Zhu, M.T. Bedford, D.M. Glover, and M. Zernicka-Goetz. 2018. CARM1 and Paraspeckles Regulate Pre-implantation Mouse Embryo Development. Cell. 175:1902–1916.e13. doi:10.1016/j.cell.2018.11.027.

Jedrusik, A., D.E. Parfitt, G. Guo, M. Skamagki, J.B. Grabarek, M.H. Johnson, P. Robson, and M. Zernicka-Goetz. 2008. Role of Cdx2 and cell polarity in cell allocation and specification of trophectoderm and inner cell mass in the mouse embryo. Genes Dev. 22:2692–2706. doi:10.1101/gad.486108.

Johnson, M.H., B. Maro, and M. Takeichi. 1986. The role of cell adhesion in the synchronization and orientation of polarization in 8-cell mouse blastomeres. J Embryol Exp Morphol. 93:239–55.

Johnson, M.H., and C.A. Ziomek. Induction of Polarity in Mouse 8-Cell Blastomeres : Specificity, Geometry, and Stability.

Korotkevich, E., R. Niwayama, A. Courtois, S. Friese, N. Berger, F. Buchholz, and T. Hiiragi. 2017. The Apical Domain Is Required and Sufficient for the First Lineage Segregation in the Mouse Embryo. Dev Cell. 40:235–247.e7. doi:10.1016/j.devcel.2017.01.006.

Koyama, H., H. Suzuki, X. Yang, S. Jiang, and R.H. Foote. 1994. Analysis of Polarity of Bovine and Rabbit Embryos by Scanning Electron Microscopy’. 50. 163–170 pp.

Laan, L., N. Pavin, J. Husson, G. Romet-Lemonne, M. van Duijn, M.P. López, R.D. Vale, F. Jülicher, S.L. Reck-Peterson, and M. Dogterom. 2012a. Cortical Dynein Controls Microtubule Dynamics to Generate Pulling Forces that Position Microtubule Asters. Cell. 148:502–514. doi:10.1016/j.cell.2012.01.007.

Laan, L., S. Roth, and M. Dogterom. 2012b. End-on microtubule-dynein interactions and pulling-based positioning of microtubule organizing centers. Cell Cycle. 11:3750–3757. doi:10.4161/cc.21753.

Larsen, E.C., O.B. Christiansen, A.M. Kolte, and N. Macklon. 2013. New insights into mechanisms behind miscarriage. BMC Med. 11:154. doi:10.1186/1741-7015-11-154.

Leung, C.Y., and M. Zernicka-Goetz. 2013. Angiomotin prevents pluripotent lineage differentiation in mouse embryos via Hippo pathway-dependent and -independent mechanisms. Nat Commun. 4:2251. doi:10.1038/ncomms3251.

Lim, H.Y.G., Y.D. Alvarez, M. Gasnier, Y. Wang, P. Tetlak, S. Bissiere, H. Wang, M. Biro, and N. Plachta. 2020. Keratins are asymmetrically inherited fate determinants in the mammalian embryo. Nature. 585:404–409. doi:10.1038/s41586-020-2647-4.

Louvet, S., J. Aghion, A. Lica Santa-Maria, P. Mangeat, and B. Maro. 1996. Ezrin Becomes Restricted to Outer Cells Following Asymmetrical Division in the Preimplantation Mouse Embryo. 177. 568–579 pp.

McDole, K., and Y. Zheng. 2012. Generation and live imaging of an endogenous Cdx2 reporter mouse line. genesis. 50:775–782. doi:10.1002/dvg.22049.

Mitsui, K., Y. Tokuzawa, H. Itoh, K. Segawa, M. Murakami, K. Takahashi, M. Maruyama, M. Maeda, and S. Yamanaka. 2003. The Homeoprotein Nanog Is Required for Maintenance of Pluripotency in Mouse Epiblast and ES Cells. Cell. 113:631–642. doi:10.1016/S0092-8674(03)00393-3.

Morris, S.A., R.T.Y. Teo, H. Li, P. Robson, D.M. Glover, and M. Zernicka-Goetz. 2010. Origin and formation of the first two distinct cell types of the inner cell mass in the mouse embryo. Proc Natl Acad Sci U S A. 107:6364–6369. doi:10.1073/pnas.0915063107.

Nishioka, N., K. ichi Inoue, K. Adachi, H. Kiyonari, M. Ota, A. Ralston, N. Yabuta, S. Hirahara, R.O. Stephenson, N. Ogonuki, R. Makita, H. Kurihara, E.M. Morin-Kensicki, H. Nojima, J. Rossant, K. Nakao, H. Niwa, and H. Sasaki. 2009. The Hippo Signaling Pathway Components Lats and Yap Pattern Tead4 Activity to Distinguish Mouse Trophectoderm from Inner Cell Mass. Dev Cell. 16:398–410. doi:10.1016/j.devcel.2009.02.003.

Nishioka, N., S. Yamamoto, H. Kiyonari, H. Sato, A. Sawada, M. Ota, K. Nakao, and H. Sasaki. 2008. Tead4 is required for specification of trophectoderm in pre-implantation mouse embryos. Mech Dev. 125:270–283. doi:10.1016/j.mod.2007.11.002.

Niwa, H., J.-I. Miyazaki, and A.G. Smith. 2000. Quantitative expression of Oct-3/4 defines differentiation, dedifferentiation or self-renewal of ES cells.

Niwayama, R., P. Moghe, Y.J. Liu, D. Fabrèges, F. Buchholz, M. Piel, and T. Hiiragi. 2019. A Tug-of-War between Cell Shape and Polarity Controls Division Orientation to Ensure Robust Patterning in the Mouse Blastocyst. Dev Cell. 51:564–574.e6. doi:10.1016/j.devcel.2019.10.012.

Panamarova, M., A. Cox, K.B. Wicher, R. Butler, N. Bulgakova, S. Jeon, B. Rosen, R.H. Seong, W. Skarnes, G. Crabtree, and M. Zernicka-Goetz. 2016. The BAF chromatin remodelling complex is an epigenetic regulator of lineage specification in the early mouse embryo. Development (Cambridge). 143:1271–1283. doi:10.1242/dev.131961.

Parfitt, D.-E., and M. Zernicka-Goetz. 2010. Epigenetic Modification Affecting Expression of Cell Polarity and Cell Fate Genes to Regulate Lineage Specification in the Early Mouse Embryo. Mol Biol Cell. 21:2649–2660. doi:10.1091/mbc.E10.

Piotrowska-Nitsche, K., A. Perea-Gomez, S. Haraguchi, and M. Zernicka-Goetz. 2005. Four-cell stage mouse blastomeres have different developmental properties. Development. 132:479–490. doi:10.1242/dev.01602.

Plachta, N., T. Bollenbach, S. Pease, S.E. Fraser, and P. Pantazis. 2011. Oct4 kinetics predict cell lineage patterning in the early mammalian embryo. Nat Cell Biol. 13:117–123. doi:10.1038/ncb2154.

Plusa, B., S. Frankenberg, A. Chalmers, A.K. Hadjantonakis, C.A. Moore, N. Papalopulu, V.E. Papaioannou, D.M. Glover, and M. Zernicka-Goetz. 2005. Downregulation of Par3 and aPKC function directs cells towards the ICM in the preimplantation mouse embryo. J Cell Sci. 118:505–515. doi:10.1242/jcs.01666.

Pomp, O., H.Y.G. Lim, R.M. Skory, A.A. Moverley, P. Tetlak, S. Bissiere, and N. Plachta. 2022. A monoastral mitotic spindle determines lineage fate and position in the mouse embryo. Nat Cell Biol. 24:155–167. doi:10.1038/s41556-021-00826-3.

Ralston, A., and J. Rossant. 2008. Cdx2 acts downstream of cell polarization to cell-autonomously promote trophectoderm fate in the early mouse embryo. Dev Biol. 313:614–629. doi:10.1016/j.ydbio.2007.10.054.

Reeve, W.J.D., and C.A. Ziomek. 1981. Distribution of microvilli on dissociated blastomeres from mouse embryos: evidence for surface polarization at compaction. 62. 339–350 pp.

Schaniel, C., Y.-S. Ang, K. Ratnakumar, C. Cormier, T. James, E. Bernstein, I.R. Lemischka, and P.J. Paddison. 2009. Smarcc1/Baf155 Couples Self-Renewal Gene Repression with Changes in Chromatin Structure in Mouse Embryonic Stem Cells. Stem Cells. 27:2979–2991. doi:10.1002/stem.223.

Schindelin, J., I. Arganda-Carreras, E. Frise, V. Kaynig, M. Longair, T. Pietzsch, S. Preibisch, C. Rueden, S. Saalfeld, B. Schmid, J.-Y. Tinevez, D.J. White, V. Hartenstein, K. Eliceiri, P. Tomancak, and A. Cardona. 2012. Fiji: an open-source platform for biological-image analysis. Nat Methods. 9:676– 682. doi:10.1038/nmeth.2019.

Shen, C., A. Lamba, M. Zhu, R. Zhang, M. Zernicka-Goetz, and C. Yang. 2022. Stain-free detection of embryo polarization using deep learning. Sci Rep. 12. doi:10.1038/s41598-022-05990-6.

Siracusa, G., D.G. Whittingham, and M. de Felici. 1980. The effect of microtubule- and microfilament-disrupting drugs on preimplantation mouse embryos. Development. 60:71–82. doi:10.1242/dev.60.1.71.

Skamagki, M., K.B. Wicher, A. Jedrusik, S. Ganguly, and M. Zernicka-Goetz. 2013. Asymmetric Localization of Cdx2 mRNA during the First Cell-Fate Decision in Early Mouse Development. Cell Rep. 3:442–457. doi:10.1016/j.celrep.2013.01.006.

Strumpf, D., C.A. Mao, Y. Yamanaka, A. Ralston, K. Chawengsaksophak, F. Beck, and J. Rossant. 2005. Cdx2 is required for correct cell fate specification and differentiation of trophectoderm in the mouse blastocyst. Development. 132:2093–2102. doi:10.1242/dev.01801.

Sutherland, A., and P.G. Calarco-Gillam. 1983. Analysis of Compaction in the Preimplantation Mouse Embryo. 100. 328–338 pp.

Tabansky, I., A. Lenarcic, R.W. Draft, K. Loulier, D.B. Keskin, J. Rosains, J. Rivera-Feliciano, J.W. Lichtman, J. Livet, J.N.H. Stern, J.R. Sanes, and K. Eggan. 2013. Developmental bias in cleavage-stage mouse blastomeres. Current Biology. 23:21–31. doi:10.1016/j.cub.2012.10.054.

Torres-Padilla, M.E., D.E. Parfitt, T. Kouzarides, and M. Zernicka-Goetz. 2007. Histone arginine methylation regulates pluripotency in the early mouse embryo. Nature. 445:214–218. doi:10.1038/nature05458.

Vinot, S., T. Le, S. Ohno, T. Pawson, B. Maro, and S. Louvet-Vallée. 2005. Asymmetric distribution of PAR proteins in the mouse embryo begins at the 8-cell stage during compaction. Dev Biol. 282:307–319. doi:10.1016/j.ydbio.2005.03.001.

White, M.D., J.F. Angiolini, Y.D. Alvarez, G. Kaur, Z.W. Zhao, E. Mocskos, L. Bruno, S. Bissiere, V. Levi, and N. Plachta. 2016. Long-Lived Binding of Sox2 to DNA Predicts Cell Fate in the Four-Cell Mouse Embryo. Cell. 165:75–87. doi:10.1016/j.cell.2016.02.032.

Zernicka-Goetz, M., J. Pines, S. Hunter McLean, J.P. Dixon, K.R. Siemering, J. Haseloff, and M.J. Evans. 1997. Following cell fate in the living mouse embryo. Development. 124:1133–1137. doi:10.1242/dev.124.6.1133.

Zhu, M., J. Cornwall-Scoones, P. Wang, C.E. Handford, J. Na, M. Thomson, and M. Zernicka-Goetz. 2020. Developmental clock and mechanism of de novo polarization of the mouse embryo. Science (1979). 370. doi:10.1126/science.abd2703.

Zhu, M., C.Y. Leung, M.N. Shahbazi, and M. Zernicka-Goetz. 2017. Actomyosin polarisation through PLC-PKC triggers symmetry breaking of the mouse embryo. Nat Commun. 8. doi:10.1038/s41467-017-00977-8.

Zhu, M., M. Shahbazi, A. Martin, C. Zhang, B. Sozen, R.S. Mandelbaum, R.J. Paulson, M.A. Mole, M. Esbert, R.T. Scott, A. Campbell, S. Fishel, V. Gradinaru, H. Zhao, K. Wu, Z.-J. Chen, E. Seli, M.J. De Los Santos, and M. Zernicka Goetz. 2021. Human embryo polarization requires PLC signaling to mediate trophectoderm specification. doi:10.7554/eLife.

Ziomek C.A., and M.H. Johnson. 1982. The Roles of Phenotype and Position in Guiding the Fate of 16-Cell Mouse Blastomeres. 91. 440–447 pp.

